# The Arabidopsis F-box protein FBW2 degrades AGO1 to avoid spurious loading of illegitimate small RNA

**DOI:** 10.1101/2021.03.24.436811

**Authors:** Thibaut Hacquard, Marion Clavel, Patricia Baldrich, Esther Lechner, Imma Pérez-Salamó, Mikhail Schepetilnikov, Benoît Derrien, Marieke Dubois, Philippe Hammann, Lauriane Kuhn, Danaé Brun, Nathalie Bouteiller, Hervé Vaucheret, Blake C. Meyers, Pascal Genschik

**Affiliations:** Institut de biologie moléculaire des plantes, CNRS, Université de Strasbourg, Strasbourg, France; Gregor Mendel Institute, Austrian Academy of Sciences, Vienna, Austria; Donald Danforth Plant Science Center, Saint Louis, Missouri (USA); DNA SCRIPT, Paris, 75014, France; VIB-UGent Center for Plant Systems Biology, Ghent University, Belgium; Plateforme Protéomique Strasbourg Esplanade du CNRS, Université de Strasbourg, Strasbourg, France; Institut Jean-Pierre Bourgin, INRAE, AgroParisTech, Université Paris-Saclay, 78000, Versailles, France; Division of Plant Sciences, University of Missouri, Columbia, Missouri, USA

## Abstract

RNA silencing is a conserved mechanism in eukaryotes and is involved in development, heterochromatin maintenance and defense against viruses. In plants, ARGONAUTE1 (AGO1) protein plays a central role in both microRNA (miRNA) and small interfering RNA (siRNA)-directed silencing and its expression is regulated at multiple levels. Here, we report that the F-box protein FBW2 targets proteolysis of AGO1 by a CDC48-mediated mechanism. We found that FBW2 assembles an SCF complex that recognizes the MID-PIWI domain of AGO1 and requires its C-terminal domain containing a GW motif for AGO1 turnover. We showed that FBW2 prefers the unloaded and some mutated forms of AGO1 protein. While *FBW2* loss of function does not lead to strong growth or developmental defects, it significantly increases RNA silencing activity. Interestingly, under conditions in which small RNA production or accumulation is affected, the failure to degrade AGO1 in *fbw2* mutants becomes more deleterious for the plant. Hence, the non-degradable AGO1 protein assembles high molecular weight complexes and binds illegitimate small RNA leading to the cleavage of new target genes that belong to stress responses and cellular metabolic processes. Therefore, the control of AGO1 homeostasis by ubiquitin ligases plays an important role in quality control to avoid off-target cleavage.

## Introduction

In eukaryotes, RNA silencing is crucial for development and plays major roles in response to the environment, including pathogens, as well as in epigenetic control of transposable elements. This pathway involves processing of double-stranded (ds)RNA by the RNase III enzyme Dicer, into small RNA of 21-to-24 nucleotides in length (Ghildiyal and Zamore, 2009). These small RNA are known to associate with ARGONAUTE (AGO) proteins to form RNA-induced silencing complexes (RISCs (Meister, 2013; Poulsen et al., 2013). RISCs are programmed by the bound small RNA to specifically interact with transcripts based on sequence complementarity, resulting in their down-regulation. Plant small RNA fall into two broad categories (Axtell, 2013). The first consists of microRNA, that are excised from stem-loop structures arising from non-coding *MIR* genes and act by post-transcriptionally repressing the levels of mRNA to which they are partly complementary. The second category encompasses so-called siRNA, which are processed from long double stranded RNA arising from a variety of sources (transposons, endogenous inverted repeats, viral RNA, transgenes) and act as repressor of expression, either transcriptionally or post-transcriptionally by mediating RNA cleavage and/or translational repression.

The *Arabidopsis thaliana* (hereafter referred to as Arabidopsis) genome encodes 10 ARGONAUTE paralogues (Vaucheret, 2008) that all have a similar domain organization and ability to bind to small RNA, although the nature and sequence of the small RNA bound by different AGOs varies greatly. Both genetic and biochemical analyses have revealed that AGO1 plays a central role in both miRNA and siRNA-directed silencing (Mi et al., 2008). Hence AGO1 loaded with miRNA mediates endonucleolytic cleavage of target transcripts (Baumberger and Baulcombe, 2005), but a fraction of transcripts can also undergo repression of protein translation (Brodersen et al., 2008; Li et al., 2013). By its ability to bind certain virus-derived siRNA, AGO1 is also an important player in plant antiviral silencing (Morel et al., 2002; Azevedo et al., 2010). Thus, upon infection by an RNA virus, dsRNA produced by intramolecular RNA folding or from replication intermediates is processed into virus-derived siRNA (vsiRNA), that are loaded into AGO1 (and other AGOs) to direct the silencing of the complementary viral RNA.

Previous work revealed that Viral Suppressor of RNA silencing (VSR) proteins P0 from poleroviruses encode F-box proteins that hijack the host S phase kinase-associated protein1 (SKP1)-cullin 1 (CUL1)-F-box protein (SCF) ubiquitin-protein ligase (E3) to promote AGO1 degradation (Pazhouhandeh et al., 2006; Baumberger et al., 2007; Bortolamiol et al., 2007; Csorba et al., 2010). It was subsequently shown that P0 triggers the vacuolar degradation of membrane-bound AGO1 via an autophagy-related process (Derrien et al., 2012) (Michaeli et al., 2019). Note that VSRs of other viruses can also promote AGO1 degradation by a different pathway involving the proteasome (Chiu et al., 2010). Beyond manipulations of AGO1 turnover by VSRs, still little is known about post-translational regulations of AGO1 in a non-viral context. Nonetheless, it was shown that mutations affecting miRNA biogenesis and/or accumulation and thus disturbing RISC assembly, result in AGO1 turnover, suggesting that the underlying mechanisms contribute to the normal cellular homeostasis of AGO1 (Derrien et al., 2012). Moreover, this homeostatic regulation in which miRNA availability controls AGO protein stability is conserved across kingdoms as it was also observed in mammalian and *Drosophila melanogaster* cells (Martinez and Gregory, 2013; Smibert et al., 2013). How metazoan AGO proteins are degraded is also not well understood. For instance, it was shown that the inhibition of HSP90 function triggered the degradation of human unloaded Ago1 and Ago2 proteins (Johnston et al., 2010), an effect that could be alleviated, at least partially, by the proteasome inhibitor MG132. However, human Ago2 is also subjected to degradation as a miRNA-free entity by selective autophagy through a pathway involving NDP52, a known autophagy receptor, which confers cargo selectivity typically by recognizing conjugated ubiquitin (Gibbings et al., 2012). Nevertheless, for both proteasomal and autophagy-dependent degradation pathways of AGO proteins, the identity of the ubiquitin E3 ligase(s) involved remained unclear until recently. A first hint came with the identification of a RING-type E3 ubiquitin ligase from Drosophila named Iruka, which preferentially binds and ubiquitylates empty Ago1 (Kobayashi et al., 2019a). Moreover, in mammals, it was recently shown that extensive miRNA–target complementarity can trigger AGO proteasomal decay exposing miRNA for degradation (Shi et al., 2020)(Han et al., 2020).

In Arabidopsis, one candidate F-box protein that controls AGO1 protein homeostasis is FBW2. FBW2 interacts with several Skp1-like (ASK) proteins in yeast two hybrid interactions and was initially identified by a genetic suppressor screen of a null allele of *SQUINT* (*SQN*), encoding a Cyclophilin-40 chaperon acting as a positive regulator of AGO1 activity (Risseeuw et al., 2003) (Earley et al., 2010). While *fbw2* mutant plants did not show a strong increase of endogenous AGO1 protein, likely because of mir168-based feedback effect (Mallory and Vaucheret, 2010), overexpression of *FBW2* in transgenic Arabidopsis lines produced a phenotype similar to hypomorphic *ago1* mutant alleles and reduced AGO1 protein level (Earley et al., 2010). Both loss of *FBW2* and over-expression of *FBW2* affect AGO1 protein levels without affecting the *AGO1* transcript, suggesting that FBW2 regulates AGO1 post-transcriptionally. However, the molecular basis of this phenotype is still unclear, and one cannot exclude that FBW2 modulates AGO1 protein level only in an indirect way. Moreover, under standard growth conditions, *fbw2* mutant plants exhibit no visible alteration in development, thus raising the question of the physiological role of this F-box protein (Earley et al., 2010). Interestingly, it was recently shown that *FBW2* is transcriptionally repressed by CURLY LEAF (CLF), encoding a subunit of the Polycomb Repressor Complex 2 (PRC2) and that this regulation may be important for AGO1 protein homeostasis when plants are exposed to UV radiation (Ré et al., 2019).

In the present work, we investigated the molecular mechanism by which FBW2 controls the stability of AGO proteins. We showed that *in planta*, FBW2 does not degrade all AGOs equally, but preferentially degrades AGO1 through its MID-PIWI domain. FBW2 is part of an SCF complex that interacts directly with AGO1 to trigger its degradation by a CDC48-mediated mechanism. Our results indicate that FBW2 plays a critical role to maintain AGO1 proteostasis by preferentially degrading its unloaded form. However, FBW2 association with membranous fractions suggests that it may also interact with AGO1 when it is loaded. Interestingly, in mutant plants lacking *FBW2* and which are impaired in small RNA accumulation, stabilized AGO1 further worsens their phenotype. Hence, we show that the non-degradable AGO1 protein assembles high molecular complexes and binds illegitimate small RNA leading to the cleavage of new target genes. Our studies identify a mechanism to avoid AGO1 spurious loading of small RNA, which could conditionally become detrimental for cells.

## Results

### FBW2 degrades AGO1 through the MID-PIWI domain

To better understand how FBW2 is involved in the control of AGO protein homeostasis, we first tested its ability to degrade different AGO proteins in a transient expression assay. Arabidopsis AGO1, AGO2, AGO3, AGO4 and AGO5, representative members of the three AGO phylogenetic clades (Vaucheret, 2008), were tagged with a Flag-tag and transiently expressed in the presence of 3HA-FBW2 (3x Human influenza hemagglutinin (HA)-epitope at the N-terminus of FBW2) or GUS (as a control) in *Nicotiana benthamiana* (*N. benthamiana*) leaves, and the level of each AGO protein was assessed with the Flag antibody (Fig. 1A). In this assay, 3HA-FBW2 was able to degrade AGO1 and AGO5 and to a lesser extent AGO2 and AGO3, but not AGO4, which was insensitive to the F-box protein. Note that AGO5 belongs to the same phylogenetic clade as AGO1, suggesting that members of this clade are better substrates for FBW2.

**Fig. 1.**
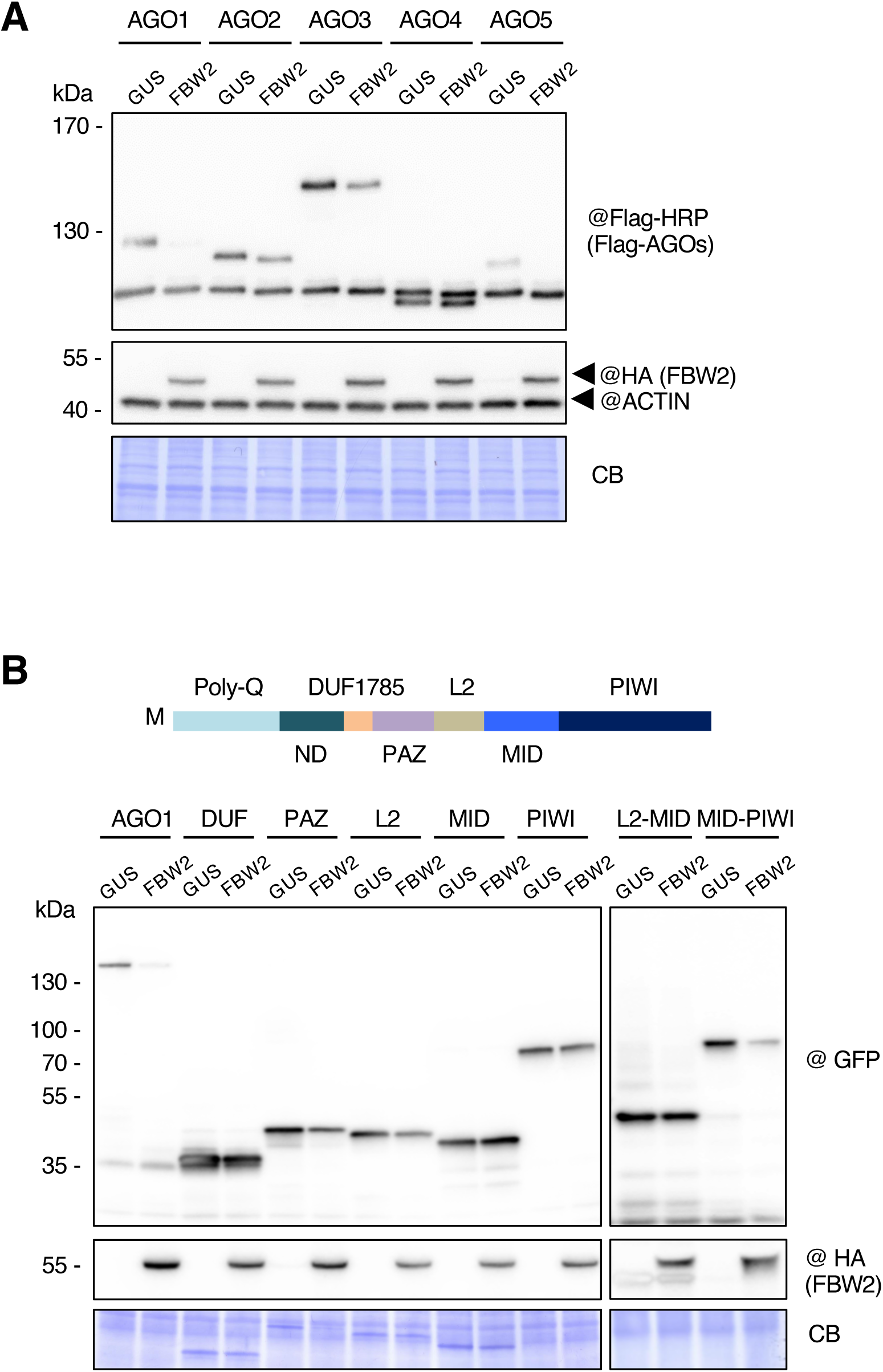
FBW2 triggers the degradation of the AGO1 Mid-Piwi domain. **(A)** FBW2 degrades effectively AGO1 and to lesser extent others AGOs. Different Arabidopsis Flag-AGO proteins, under the control of the CaMV 35S promoter, were transiently expressed in *N. benthamiana* leaves in absence (35S:GUS) or presence of FBW2 (35S:3HA-FBW2). Proteins were extracted 72 hours after agro-infiltration and AGO proteins were detected by western blot with Flag antibodies. Probing with the ACTIN antibody and Coomassie blue (CB) staining were used as loading controls. (B) FBW2 triggers the degradation of the AGO1 Mid-Piwi domain. Schematic representation of Arabidopsis AGO1 colour-coded protein domains: DUF1785: Domain of Unknown Function 1785. PAZ: Piwi-Argonaute-Zwille. L2: Linker 2. MID: Middle domain. Agrobacteria harbouring single and combined AGO1 protein domains fused to GFP, under the control of the CaMV 35S promoter, were infiltrated at an OD of 0,3 in *N. benthamiana* leaves in absence (35S:GUS) or presence of FBW2 (35S:3HA-FBW2). Proteins were extracted 72 hours after agro-infiltration and fusion protein levels were assessed by immunoblot using GFP and HA antibodies. Coomassie blue (CB) staining was used as loading control.

To address the question of which domain of AGO1 is required for the FBW2-mediated degradation, we expressed in *N. benthamiana* leaves different constructs covering each domain of AGO1 fused to GFP, in the absence or presence of 3HA-FBW2. Though we observed a partial degradation of the PAZ and L2, most of these domains taken individually were not sufficient to trigger FBW2-mediated decay (Fig. 1B). However, by combining the MID and PIWI domains, covering the C-terminal part of AGO1, the fusion protein was degraded by FBW2 as efficiently as the full-length AGO1 protein (Fig. 1B; Fig. S1).

Next, we aimed to further investigate this mechanism in stable Arabidopsis lines overexpressing tagged 3HA-FBW2. For this purpose, we first verified in *N. benthamiana* leaves that the tagged protein was as efficient as the native F-box to degrade AGO1 (Fig. S2A). We then generated 35S:3HA-FBW2 stable transgenic lines (hereafter named FBW2OE lines) and we observed that out of 50 lines analysed, none of them showed severe developmental defects that would be expected for a strong depletion of AGO1. Nevertheless, we could identify several lines in which 3HA-FBW2 protein was overexpressed (Fig. S2B) and we continued with one of them (line #10). In this line, we observed a significant decrease in AGO1 protein despite a higher level of its transcript (Fig. 2A). A phenotypic examination of the FBW2OE line showed a reduction of leaf growth and an increase in the number of lateral roots, whereas the *fbw2-4* knockout line did not produce a visible phenotype under standard growth conditions (Fig. S2C-D).

**Fig. 2.**
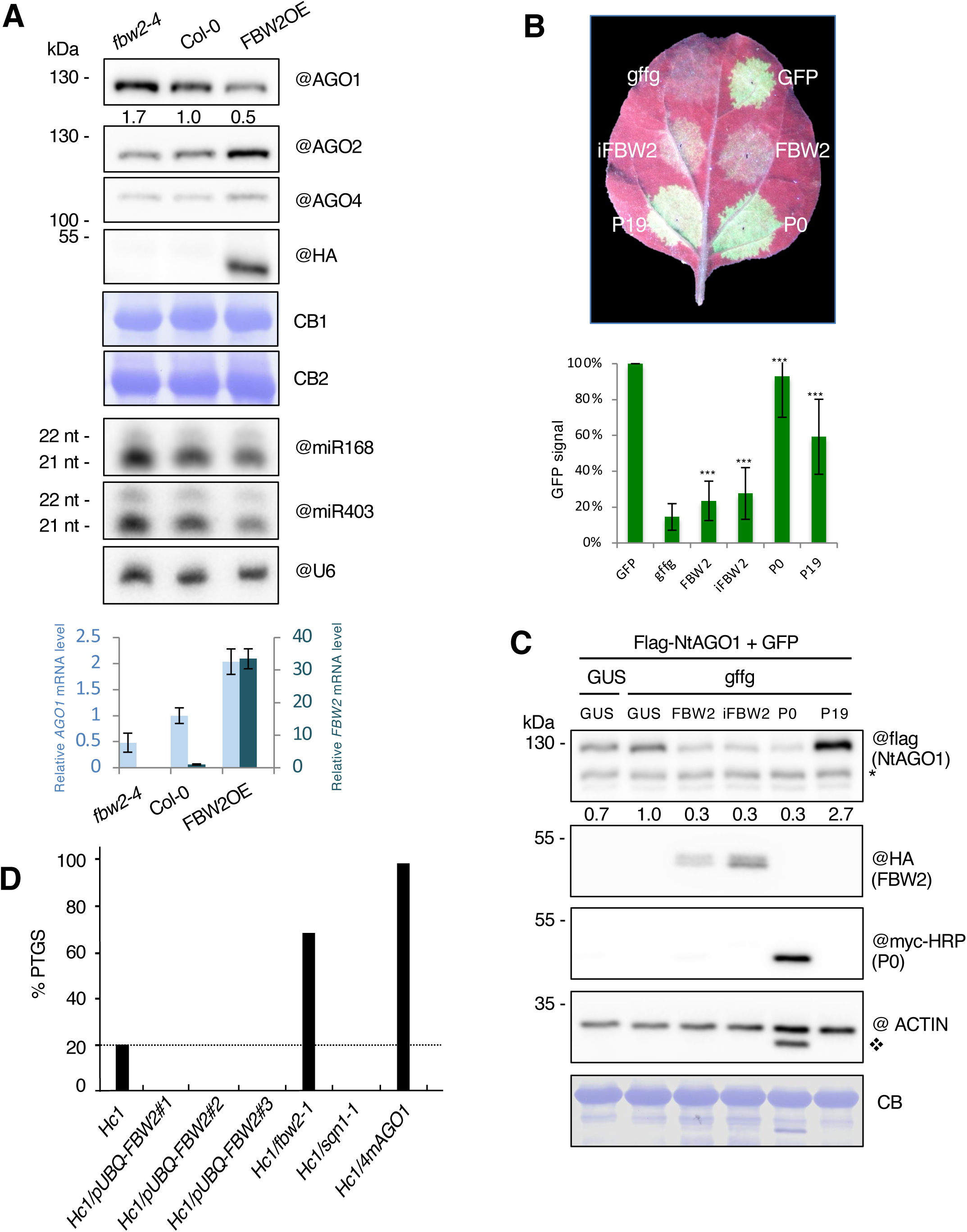
Function of FBW2 in RNA silencing. **(A)** FBW2 overexpression leads to partial degradation of AGO1. Top panel: Immunoblot analysis of AGO1, AGO2 and AGO4 protein contents in the *fbw2-4* mutant allele, in Col-O and in FBW2OE line (35S:3HA-FBW2 line 10). Seedlings grown on MS medium were harvested at 8 days and protein extracts were analysed by immunoblotting of AGOs using specific antibodies and of FBW2 using the HA antibody. Coomassie blue (CB) staining was used as loading controls (CB1 corresponds to AGO1 and CB2 to 3HA-FBW2). AGO1 signal was quantified by ImageJ, normalized to the corresponding CB. Numbers are indicated below the panel as relative to Col-0 set at 1.0. Middle panel: Small RNA gel blot analysis of the steady state accumulation of the indicated miRNAs taken from the same material as above. U6 RNA level was used as a loading control. Bottom panel: RT-qPCR analysis of *AGO1* and *FBW2* transcript levels in the *fbw2-4*, Col-O and FBW2OE line. Total RNA samples were extracted from the same material as above. **(B)** FBW2 is a weak endogenous suppressor of RNA silencing. Upper panel, picture of a *N. benthamiana* leaf 72 hours after infiltration with agrobacteria harbouring a 35S:Flag-NtAGO1 and a 35S:GFP construct plus either the following constructs: 35S:GUS or 35S:GFFG (GFP mRNA hairpin) together with either 35S:3HA-FBW2 (only the coding sequence), 35S:3HA-iFBW2 (coding sequence including an intron), 35S:P0-6myc or 35S:P19. Bottom panel, the intensity of GFP signal in the infiltration area was measured with an Ettan DIGE imager (GE healthcare) and normalized to the GFP control condition. *** p < 0,001 (T-test) as compared to *GFFG*. **(C)** Western blot of protein extracts from tissues sampled 72 hours after agro-infiltration (shown in (B). Coomassie blue (CB) staining and ACTIN protein level were used as a loading control. The “@” symbol indicates hybridization with the corresponding antibodies. NtAGO1 signal was quantified by ImageJ, normalized to the corresponding CB. Numbers are indicated below the panel as relative to the control set at 1.0. * Non-specific band; v Remaining signal from P0-6myc hybridization. **(D)** Modulating FBW2 level impacts transgene S-PTGS efficiency. GUS activity was measured in leaves of 8 week-old plants of the indicated genotypes. For each genotype, 96 plants were analysed. S-PTGS efficiency is expressed as the percentage of plants exhibiting GUS activity below 50 FLUO.min^-1^.ug^-1^.

In parallel, we also generated Arabidopsis transgenic lines in which 3HA-FBW2 was expressed under the control of the β-Estradiol (β-Es)-inducible promoter XVE (Zuo et al., 2000). Out of 65 stable lines, we selected seven XVE:3HA-FBW2 lines for high expression of 3HA-FBW2 upon β-Es induction without leakage. By monitoring FBW2 and AGO1 protein levels in a kinetic experiment upon β-Es treatment for 12 days, we noticed that a transient expression of 3HA-FBW2 was not sufficient to efficiently degrade AGO1 (Fig. S3A). This situation contrasts with the viral encoded F-box protein P0, which significantly affects AGO1 protein level already at 5 days of induction (Fig. S3B). Next, we introduced by crossing the XVE:3HA-FBW2 construct in different *ago1* mutant backgrounds, including *ago1-57, ago1-27* and *ago1-38* (Fig. S3C). Of particular interest was the *ago1-57* mutation in the DUF1785, which abrogates SCF-dependent P0 interaction with AGO1 (Derrien et al., 2018), as we wondered whether this mutation would also affect FBW2-mediated AGO1 degradation. However, we observed that AGO1-57 protein was degraded by FBW2 (Fig. S3C and S3D), indicating that the AGO1 degron recognised by this F-box protein is not located in the DUF1785, which is consistent with the transient degradation assays (Fig. S1). In contrast to 3HA-FBW2-induction in Col-0, for which FBW2 protein was well induced at the start of the treatment but quickly disappeared over time (Fig. S3A), in the *ago1-57* mutant background, the FBW2 protein accumulated for a longer period of time leading to a more pronounced AGO1 protein destabilization (Fig. S3D, upper panel). qRT-PCR analyses showed that the transcript level of *3HA-FBW2* decreased less over time in *ago1-57* background if compared to Col-0 (Fig. S3D, bottom panel), suggesting the suppression of the transgene silencing by the mutation. Similar results were also obtained with a mutant of *RNA-dependent RNA polymerase 6* (*SGS2/SDE1/RDR6)* (Mourrain et al., 2000) affecting sense transgene-mediated post-transcriptional gene silencing (S-PTGS) leading to a higher degradation rate of AGO1, likely as a consequence of an increased and more sustained level of *FBW2* expression (Fig. S3E). It is noteworthy that among different AGO1 mutations, the AGO1-27 protein was the most susceptible to FBW2-mediated degradation, which cannot be solely explained by increased FBW2 expression (Fig. S3C) and this was also observed when *FBW2* was constitutively overexpressed in this mutant background (see below).

To address the question of the specificity of FBW2 towards other Arabidopsis AGO proteins, we monitored the protein levels of AGO2 and AGO4 in our FBW2 overexpressor lines. In contrast to the effect of *FBW2* on the steady state level of AGO1, both AGO2 and AGO4 were insensitive to the degradation activity of the F-box protein (Fig. 2A; (Fig. S3C-E). Instead, AGO2 protein levels were even increased when FBW2 was overexpressed and this might be attributed to the partial degradation of AGO1, which in association with miR403 targets AGO2 transcript (Allen et al., 2005). Hence, the levels of both miR168 and miR403 were decreased in FBW2OE seedlings and *AGO1* transcript level was upregulated (Fig. 2A, middle and bottom panels). Conversely, in the *fbw2-4* null mutant, AGO1 protein steady state level was slightly increased, indicating that FBW2 contributes to maintain AGO1 protein homeostasis under normal growing conditions (Fig. 2A).

### Modulating FBW2 level impacts transgene S-PTGS efficiency

As FBW2 destabilizes AGO1, we investigated its activity in suppressing RNA silencing. At first, the effect of FBW2 on inverted-repeat (IR)-PTGS was tested. For this, we used a patch assay in which a *GFP* transgene is transiently expressed in *N. benthamiana* and its silencing is triggered by a *GFFG* inverted-repeat RNA (Himber et al., 2003). In this assay, *Nicotiana tabacum* AGO1 N-terminally fused to a Flag-tag was co-expressed to monitor its protein level. As expected, when the patches were infiltrated with two strong VSRs, P0 and P19 (Csorba et al., 2015), a bright-green GFP fluorescence signal was detected (Fig. 2B, upper panel). In contrast to these VSRs, co-expression of FBW2 had only a weak impact on IR-PTGS triggered by the *GFFG* RNA. Quantification of the GFP fluorescence showed ± 10% increased fluorescence of the silenced *GFP* when FBW2 is co-expressed (Fig. 2B, bottom panel), indicating that FBW2 acts as a weak suppressor of IR-PTGS in this system. On western blot (Fig. 2C), tobacco AGO1 is only partially destabilized by FBW2, suggesting that together with other AGOs, the amount of active AGO1 in these plants is sufficient to execute IR-PTGS.

Next, the effect of FBW2 on sense (S)-PTGS was tested because S-PTGS is more sensitive to small perturbation in AGO1 activity than IR-PTGS. Indeed, the hypomoporphic *ago1-27* mutation totally impaired S-PTGS while it only decreased IR-PTGS (Parent et al., 2015). Therefore, we crossed the Arabidopsis *fbw2-1* mutant with the p35S:GUS line Hc1 (Elmayan et al., 1998). This line triggers S-PTGS in only 20% of the population at each generation, thus representing a valuable sensor to precisely monitor changes in silencing efficiency. While as expected 20% of silencing was reached in the Col-0 background, we observed that 66% of the *fbw2-1* plants were silenced (n=96; Fig. 2D). This increase in S-PTGS is consistent with the slight increase in AGO1 protein level observed in *fbw2* loss-of-function mutants (Fig. 2A). It is also consistent with the fact that a higher increase in AGO1 protein level is necessary to elevate Hc1 S-PTGS frequency up to 100%, for example in 4mAGO1 plants, which express a miR168-resistant form of the AGO1 mRNA (Fig. 2D; (Martínez De Alba et al., 2011)). We also generated three independent Hc1 lines overexpressing FBW2. To avoid potential interference between p35S:GUS and p35S:FBW2 transgenes, we used RFP-tagged FBW2 expressed under the control of the pUBQ10 promoter (Grefen et al., 2010). Analysis of the HC1/pUBQ-FBW2 lines showed a clear inhibition of S-PTGS Fig. 2D), which is an opposite phenotype to the *fbw2-1* mutation, and which mimics the effect of the *sqn-1* mutation (Fig. 2D). This result therefore confirms the previous observation that FBW2 overexpression decreased S-PTGS of the p35S:GUS line L1 (Earley et al., 2010)

### FBW2 assembles an SCF complex that interacts with AGO1

Although our data suggest that FBW2 directly and specifically targets AGO1 for degradation, this still needs to be demonstrated at the molecular level. Thus, we investigated whether FBW2 is able to interact with AGO1 *in planta*. We first examined the subcellular localization of both proteins. The coding sequence of FBW2 was fused to the Venus fluorescent protein at its N-terminus, and put under the control of its own promoter (pFBW2:Venus-FBW2). This construct was transiently co-expressed with Cyan Fluorescent Protein (CFP)-AGO1 in *N. benthamiana* leaves. Confocal imaging revealed the co-localization of both proteins in the cytosol (Fig. 3A). Note that the Venus-FBW2 protein was functional as it caused the degradation of CFP-AGO1 in this assay (Fig. 3A). Next, we immunoprecipitated (IP) 3HA-FBW2 from *Arabidopsis* plants and could show that the F-box was able to efficiently pull-down endogenous AGO1 in the presence of MLN4924 (Fig. 3B), a drug that inhibits CULLIN1 (CUL1) neddylation (Hakenjos et al., 2011). Moreover, all components of the SCF (CUL1, ASK1 and RBX1) were also pulled-down in the IP, indicating that FBW2 forms an SCF-type ubiquitin E3 ligase complex *in planta*. Accordingly, deletion or mutation of the F-box domain in FBW2 abrogated its capacity to degrade AGO1 when transiently expressed in *N. benthamiana* leaves (Fig. 3C). To further identify the interaction network of FBW2, we immunoprecipitated the F-box protein when expressed in Col-0 and in the *ago1-27* mutant, as this background showed a higher degradation rate of AGO1, and performed mass spectrometry analysis. Note that cellular extracts were crosslinked with formaldehyde before immunoprecipitation (IP). We compared 8 samples (4 samples of FBW2OE and 4 samples of FBW2OE/*ago1.27*) from two independent biological replicates to 7 control samples. Proteins significantly enriched in the FBW2 IP were highlighted by a statistical analysis, calculating normalized fold changes and adjusted p-values (Fig. 3D; Table S1). As expected, proteins of the SCF complex are predominantly enriched, such as CUL1, the CUL-like protein1, RUB1, as well as ASK1, ASK20, ASK21 and the FBW2 target AGO1. Note that AGO5 and AGO10, belonging to the same phylogenetic clade as AGO1, were also found enriched in the IP. Interestingly, a significant group of proteins co-purifying with FBW2 consists of molecular chaperones, including heat shock proteins (mainly Hsp70, Hsp80 and Hsp90) and DNAJ homologues J2/J3, for which functions of some have previously been linked to AGO1 (Iki et al., 2012)(Iki et al., 2010)(Earley and Poethig, 2011)(Sjögren et al., 2018). Strikingly, we also identified most proteasomal subunits pointing to an important function of this pathway for FBW2 and/or AGO1 proteolysis.

**Fig. 3.**
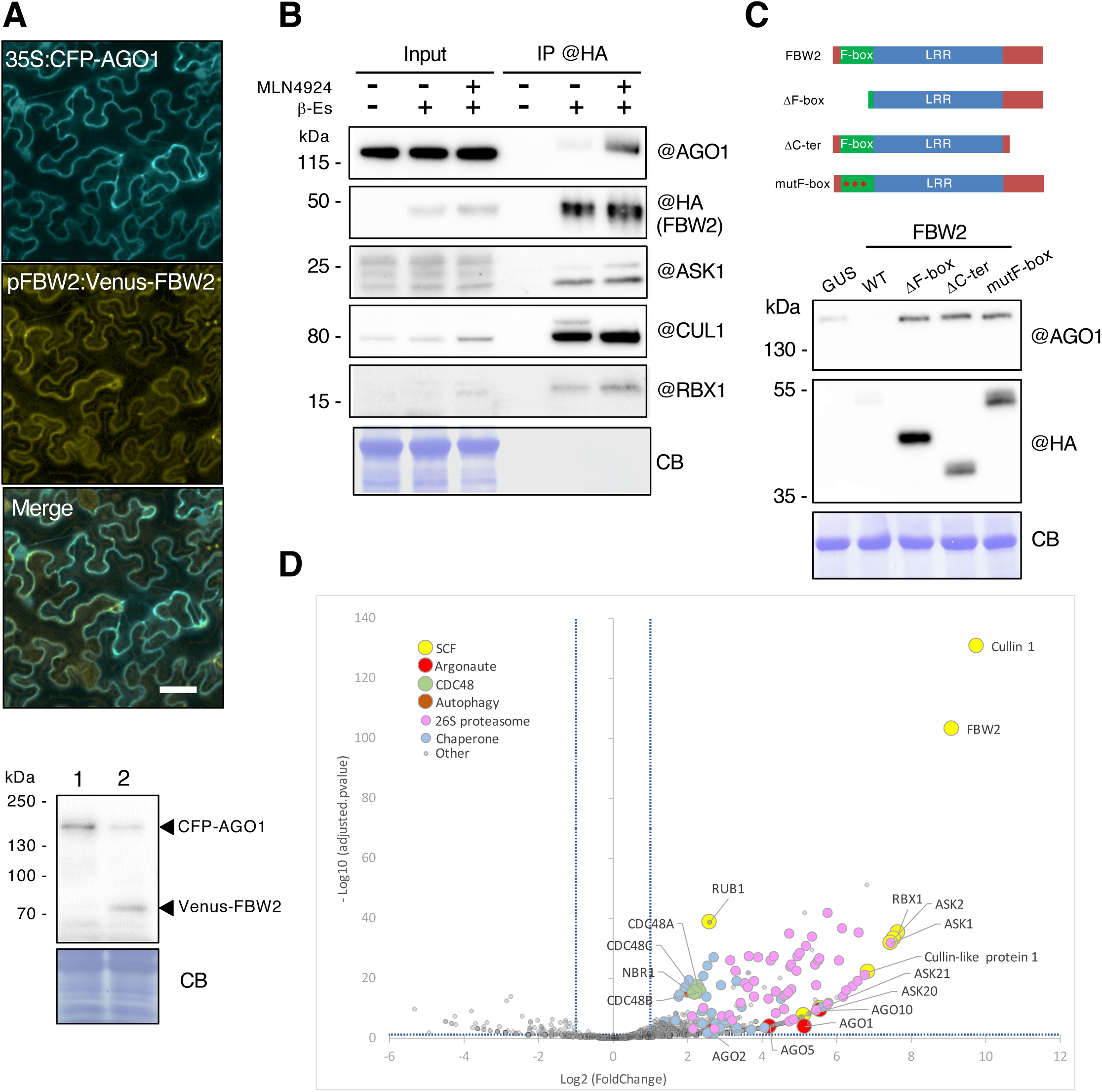
FBW2 assembles an SCF complex and directly interacts with AGO1. **(A)** Subcellular localization of CFP-AGO1 and Venus-FBW2 by confocal microscopy. Co-infiltration of 4 week-old *N. benthamiana* leaves with Agrobacteria harbouring binary vectors for the expression of fluorescent-tagged protein constructs. Bacteria were infiltrated at an OD of 0,1. Pictures were taken and tissues were sampled 3 days later. For confocal microscopy imaging, CFP and Venus were excited at 458 and 514 nm, respectively. Emission signals were recovered between 465 and 510 nm for the CFP and 520 and 596 nm for the Venus (Scale bars: 40μm). Immunodetection using GFP antibodies of protein extracts from agro-infiltrated leaves with 35S:CFP-AGO1 and pFBW2:Venus-FBW2 (lane 2) constructs is included. Expression of GUS (lane 1) serves as negative control. Coomassie blue (CB) staining was used as a loading control. **(B)** FBW2 assembles an SCF complex and interacts *in planta* with AGO1. Western blot of protein extracts from 10 day-old XVE:3HA-FBW2 seedlings. 3HA-FBW2 was immunoprecipitated with anti-HA antibodies after an overnight induction of expression in liquid MS medium supplemented with DMSO (-) or β-Es (10µM) (+). 3HA-FBW2 co-immunoprecipitates with SCF components: ASK1, CUL1 and RBX1. Blocking the SCF activity with the drug MLN4924 further allows co-immunoprecipitation of AGO1. The “@” symbol indicates hybridization with the corresponding antibodies. **(C)** Transient degradation assays of AGO1 with mutated FBW2. Western blot of protein extracts from 4 week-old *N. benthamiana* agro-infiltrated leaves. Agrobacteria harbouring a 35S:CFP-AGO1 and a 35S:3HA-FWB2 (wild type (WT), mutant and deletion) constructs were infiltrated at an OD of 0,3 and tissues were sampled 3 days later. Expression of GUS serves as control. Coomassie blue (CB) staining was used as a loading control. The “@” symbol indicates hybridization with the corresponding antibodies. The upper panel represents the schematic representation of FBW2 color-coded protein domains and deletions mutants. The mutF-box construct corresponds to 3 amino acids substitutions in the F-box domain (I21A, L25A and P26A). **(D)** FBW2 interactome revealed by immunoprecipitation and mass spectrometry. Volcano plot shows the enrichment of proteins co-purified with HA-tagged FBW2 bait as compared with Col-0 controls. Y- and x- axis display log values from adjusted p-values and fold changes, respectively. The horizontal dashed line indicates the threshold above which proteins are significantly enriched (adjusted p-values < 0.05). The vertical dashed lines indicate the fold change thresholds for FBW2-enriched proteins (log2 > 1) or Col-0-enriched proteins (log2 < -1). Six colour coded functional clusters are highlighted in the case of proteins enriched in the FBW2 co-IP samples. The source data is available in Supplementary Table S1.

To establish that FBW2 interacts directly with AGO1, we first performed yeast two-hybrid (Y2H) assays. Because AGO1 full-length protein was only very poorly expressed in this system, both the N-terminal (AGO1 NT-PAZ) and C-terminal (AGO1 L2-CT) halves of the protein were expressed separately to test for their interaction with FBW2 (Fig. 4A; Fig. S4). In agreement with our degradation assays showing that the MID-PIWI domain confers AGO1 degradability (Fig. 1B), FBW2 was able to interact only with the AGO1 C-terminal domain. To further test whether these proteins directly interact *in planta*, we performed bimolecular fluorescence complementation (BiFC) assays (Walter et al., 2004). As a negative control, we used the CULLIN3 receptor BPM5 that targets PP2CA involved in ABA signaling (Julian et al., 2019) and as a positive control SDE3, which has previously been shown to interact with AGO1 (Garcia et al., 2012). Thus, a clear cytosolic-localized YFP fluorescence signal could be detected when SDE3, but not BPM5, was co-expressed with AGO1 in Arabidopsis protoplasts (Fig. 4B-C). However, we observed that FBW2 did not interact with the full-length AGO1 in this assay and we therefore tested its C-terminal half containing the MID-PIWI domain. Consistent with the Y2H assays, a clear interaction with FBW2 was only observed with AGO1 L2-CT (Fig. 4B-C). This suggests that the domain of interaction with FBW2 is not well accessible in full-length AGO1 and that this interaction may depend on the conformation of the protein.

**Fig. 4.**
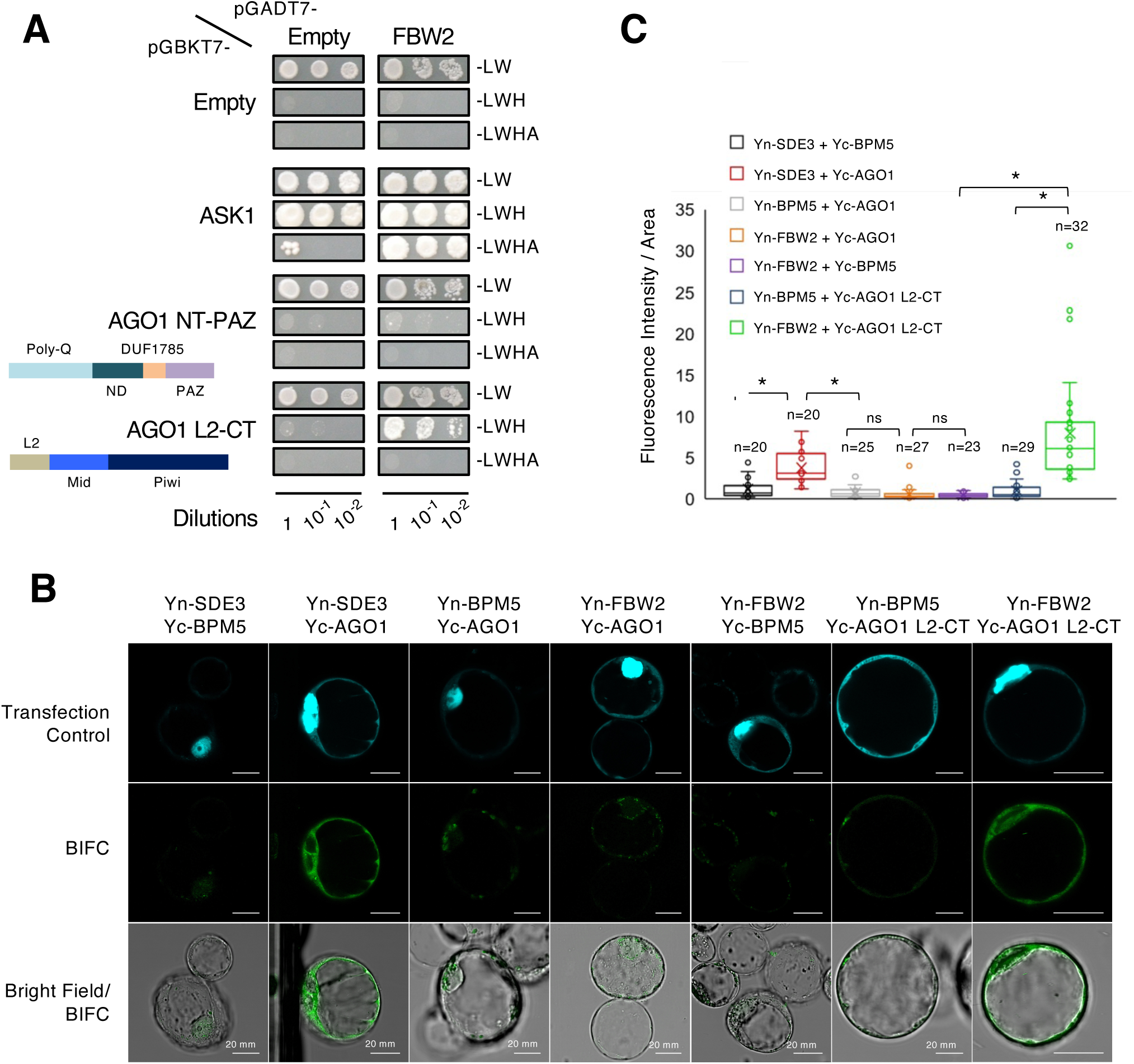
FBW2 interacts with AGO1 L2-CT in yeast and Arabidopsis protoplasts. **(A)** FBW2 interacts with the C-terminal AGO1 protein domain. Interaction between FBW2 and two AGO1 fragments (schematic representation shown on the left) were tested by yeast two-hybrid assays. ASK1 was used as positive control. Yeasts were grown for 15 days at 28°C on non-selective plates lacking leucine and tryptophan (-LW) or selective plates lacking in addition histidine (-LWH) or lacking in addition histidine and adenine (-LWHA). Note that ASK1 showed auto-activation on (-LWH), but the interaction was detected on (-LWHA) plates. (B) Pairs of constructs expressing the indicated proteins fused to the N- or C-terminal fragment of YFP (Yn or Yc) were transiently expressed in Arabidopsis cell culture protoplasts. Co-expression of SDE3, FBW2, AGO1 and AGO1-L2-CT with BPM5 protein fusions was used as negative control. The CPRF2-CFP construct was used as transfection control to identify the transformed protoplasts (Bortolamiol et al., 2007). Scale bars: 20μm. (C) Quantitative evaluation of fluorescence intensities in transformed protoplasts. Boxplots with whiskers show the median, the 25th and 75th percentiles and the interquartile ranges. Outliers are represented by dots for each construct combination. Asterisks indicate significant differences at P<0.001 among different samples by Student’s t-test. Number of transformed protoplasts used for YFP signal quantification (n) is indicated on each boxplot.

### The C-terminal domain of FBW2 is important for AGO1 degradation

Besides its F-box motif, FBW2 carries a leucine-rich repeat (LRR) domain and a C-terminal unstructured domain enriched in tryptophan (W) residues (Fig. 5). Since FBW2 principally targets AGO1 MID-PIWI domains (see above), which contains the binding pocket necessary for the interaction with GW proteins (Till et al., 2007; Elkayam et al., 2017), we wondered whether FBW2 contains an AGO-hook motif. Prediction tools (Karlowski et al., 2010) identified a single GW motif at residues 287-299. However, in addition to this stretch (including W295), we noticed that the C-terminal part of FBW2 also contains other putative GW motifs that are spaced by about 16-19 amino acids (Fig. 5B). To determine the functional importance of this domain, we expressed an FBW2 variant lacking it (Δ-Cter) in *N. benthamiana* leaves and found that the C-terminal GW domain was required for FBW2- mediated AGO1 degradation (Fig. 3C). We next site-directed mutagenized each W residue of this domain, which were replaced by alanine and we expressed the FBW2 mutated variants together with CFP-AGO1 transiently in *N. benthamiana*. While the mutation of each single W residue did not or only mildly affected AGO1 degradation, the combined mutation of four or five W residues significantly impaired it (Fig. 5B). From these results we conclude that the C-terminal domain of FBW2 is required to degrade AGO1, but its specific role in this process still remains to be established.

**Fig. 5.**
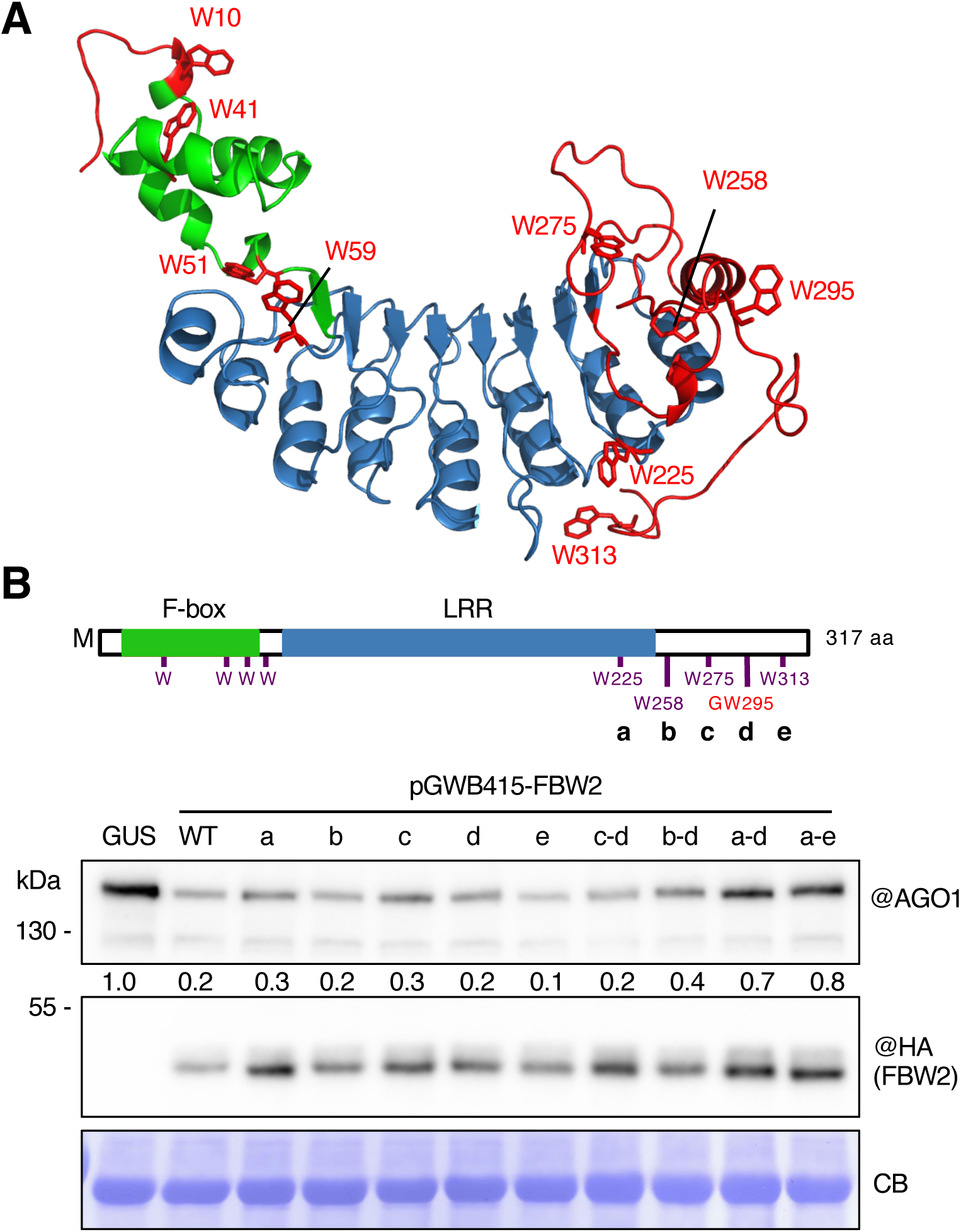
FBW2 requires its F-box and GW domain to degrade AGO1. **(A)** Prediction of FBW2 3D structure by PHYRE2 (Kelley et al., 2015). The sequence from residue 8 to 248 is structured with a high confidence (99.9%) from other known structures of F-box proteins such as Human SKP2. The F-box domain is indicated in green and the leucine-rich repeats (LRR) in blue. In red at the N-terminal and C-terminal ends are regions that could not be structured by prediction. (B) FBW2 contains a C-terminal domain enriched in tryptophan residues, which is required for AGO1 degradation. Upper panel represents FBW2 colour-coded protein domains and localization of its tryptophan (W) residues, which are either part of the F-box motif or localized in the C-terminal part of the protein. LRR: Leucine-Rich Repeats. The bottom panel shows a Western blot of protein extracts from 4 week-old *N. benthamiana* agro-infiltrated leaves. Agrobacteria harbouring CFP-AGO1 and 35S:3HA-FWB2 wild type (WT) or mutated as specified, constructs were infiltrated at an OD of 0,3 and tissues were sampled 3 days later. Expression of GUS serves as control. Coomassie blue (CB) staining was used as a loading control. The “@” symbol indicates hybridization with the corresponding antibodies. AGO1 signal was quantified by ImageJ, normalized to the corresponding CB. Numbers are indicated below the panel as relative to the control set at 1.0.

### Mechanism of FBW2-mediated AGO1 decay

To examine the mechanism by which FBW2 mediates AGO1 degradation, we first treated the FBW2OE line with the protein synthesis inhibitor cycloheximide (CHX). This revealed that the FBW2 protein rapidly decayed, with a half-life of less than 30 min (Fig. 6A). When in addition to CHX we added Bortezomib, a highly selective proteasome inhibitor (Bonvini et al., 2007), FBW2 protein level was significantly stabilized (Fig. 6B). In contrast to the unstable FBW2, which is degraded by the proteasome, the endogenous AGO1 protein level remained overall constant during the time-course of this experiment (Fig. 6A). As FBW2 was constitutively expressed under the control of the 35S promoter, it appears that the pool of AGO1 that remains insensitive to FBW2 exhibits a long half-life.

**Fig. 6.**
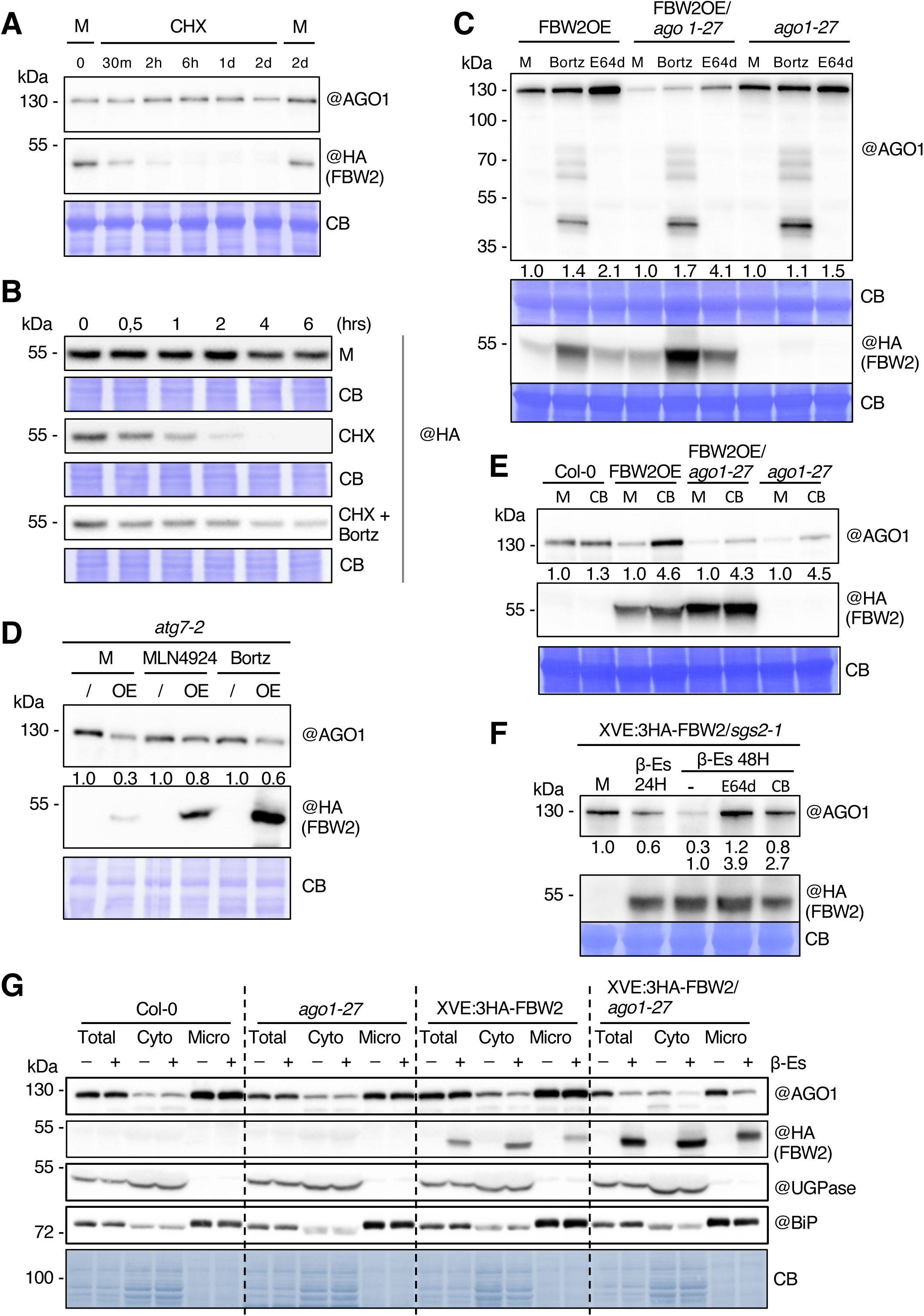
FBW2 and AGO1 protein degradation pathways. **(A)** FBW2, but not AGO1 is highly unstable. Ten day-old FBW2OE (35S:3HA-FBW2) seedlings were transferred in liquid MS medium containing either mock (M, DMSO) or the protein synthesis inhibitor cycloheximide (CHX, 100µM). At various times, the seedlings were harvested and protein extracts were analysed by immunoblotting of endogenous AGO1 and FBW2 using the indicated antibodies. Coomassie blue (CB) staining was used as a loading control. (B) Proteasome inhibition blocks FBW2 protein turnover. Ten day-old FBW2OE seedlings were transferred in liquid MS medium containing either mock (M, DMSO) or the protein synthesis inhibitor cycloheximide (CHX, 100µM) alone or CHX (100µM) combined with the proteasome inhibitor Bortezomib (Bortz, 100µM). At various times, the seedlings were harvested and protein extracts were analysed by immunoblotting of FBW2 using the HA antibody. Coomassie blue (CB) staining was used as loading controls. (C) E-64D and to a lesser extent the proteasome inhibitor Bortezomib increase AGO1 protein accumulation in FBW2OE plants. Western blot of protein extracts from 8 day-old seedlings from the indicated genotypes treated for 20 hours either with mock (M, DMSO), Bortezomib (Bortz, 100µM) or E64d (100µM). FBW2 and AGO1 proteins were detected using the indicated antibodies. Coomassie blue (CB) staining was used as loading controls. For this panel and also for panels 6D-F, AGO1 signal was quantified by ImageJ, normalized to the corresponding CB. Numbers are indicated below the panel as relative to the control set at 1.0. (D) FBW2-mediated AGO1 decay is not blocked in a core autophagy mutant. Western blot of protein extracts from 10 day-old FBW2OE/*atg7-2* seedlings. Seedlings were grown on MS medium, acclimated for 48 hours in liquid MS and subsequently treated over night with Bortezomib (Bortz, 100µM) or E64d (100µM). (**E-F**) The CDC48 inhibitor CB-5083 partially restores AGO1 protein level in FBW2 overexpressing plants. (E) Western blot of protein extracts from 8 day-old seedlings from the indicated genotypes treated for 20 hours either with mock (M, DMSO) or CB-5083 (indicated CB, 20µM). (F) Western blot of protein extracts from XVE:3HA-FBW2/*sgs2-1* seedlings treated for 24 hours with DMSO (M) or β-Es (10µM) then another 24 hours with DMSO (M) or with β-Es (10µM) or β-Es (10µM) + E64d (100µM) or β-Es (10µM) + CB-5083 (CB, 20µM). FBW2 and AGO1 proteins were detected using the indicated antibodies. Coomassie blue (CB) staining was used as loading controls. (**G**) FBW2 preferentially degrades soluble and membrane-bound compromised pool of AGO1 in the *ago1-27* background. Immunoblot analysis of the FBW2-mediated AGO1 subcellular degradation in the total protein extract (Total), cytoplasmic (Cyto) and membranous (Micro) fractions prepared from 7 day-old XVE:3HA-FBW2 β-Es-inducible lines in the background of Col-0 and *ago-1-27* grown *in vitro* on 1/2MS agar plates supplemented with 0.1% DMSO (-) or 10μM β-Es (+). Cellular fractions were probed with specific antibodies against Arabidopsis AGO1, cytoplasmic UGPase enzyme, ER luminal binding protein BiP and HA tag. Coomassie blue (CB) staining serves as a loading control. Note that 3HA-FBW2 expression level is higher in the *ago1-27* mutant background.

It was previously reported that the proteasome inhibitor MG132 was unable to restore AGO1 protein levels in *FBW2* overexpressing plants (Earley et al., 2010). To further address this question, we treated with Bortezomib Arabidopsis seedlings expressing 3HA-FBW2 in both Col-0 wild type (WT) and *ago1-27* mutant backgrounds, as the AGO1-27 protein was very sensitive to FBW2-mediated decay. Under these conditions, proteasome inhibition did only poorly block AGO1 protein degradation when *FBW2* was overexpressed (Fig. 6C). Nevertheless, we detected several AGO1 protein fragments, especially a fragment of about 48 kDa, upon Bortezomib treatment, indicating that at least a fraction and certain cleavage products of AGO1 undergo proteasome-dependent degradation. By contrast, when the lysosomal protease inhibitor E64d was used (Klionsky et al., 2016), we detected a more prominent accumulation of AGO1 in the FBW2 overexpressing lines suggesting an autophagy-related degradation pathway (Fig. 6C, F). To address this further, we crossed the FBW2OE line with the *atg7-2* knockout mutant, in which macroautophagy is genetically disrupted (Thompson et al., 2005). *ATG7* encodes an E1-like activating enzyme required to activate the ubiquitin-like proteins ATG12 and ATG8, which are essential for autophagosome biogenesis. Nevertheless, overexpression of 3HA-FBW2 in *atg7-*2 was still able to degrade AGO1, at least partially (Fig. 6D). As autophagy inhibition might be compensated by the proteasome activity (Kocaturk and Gozuacik, 2018), we treated the FBW2OE/*atg7-2* line with Bortezomib. We observed an increase in 3HA-FBW2 protein levels (supporting the efficiency of the drug treatment), but this was not the case for AGO1, which still was degraded. From these results we conclude that FBW2-mediated AGO1 decay likely employs a pathway different from canonical autophagy.

It was recently shown that the degradation of empty Ago1 protein from Drosophila requires valosin-containing protein (VCP; also known as p97 or CDC48 and hereafter called CDC48), which recognizes poly-ubiquitylated proteins via adaptor subunits (Kobayashi et al., 2019b). Interestingly, among the proteins significantly enriched in the FBW2 IP, we could identify all three Arabidopsis CDC48A-C subunits (Fig. 3D). Thus, to examine the function of these proteins in the turnover of Arabidopsis AGO1, we used CB-5083, a drug that was proven to efficiently inactivate plant CDC48 complexes (Marshall et al., 2019)(Tang et al., 2019). Remarkably, CB-5083 restored at least partially AGO1 protein levels in different genetic backgrounds regardless of whether *FBW2* was constitutively expressed or induced by β-Es (Fig. 6E-F).

Finally, we wondered whether FBW2 acts on a specific cellular pool of AGO1. Indeed, previous research has shown that AGO1 appears in membrane-free (soluble) and membrane-bound (especially associated with the endoplasmic reticulum (ER)) forms (Li et al., 2013)(Brodersen et al., 2012)(Michaeli et al., 2019). We therefore evaluated AGO1 protein level in soluble and microsomal fractions upon β-Es-inducible expression of FBW2 in WT and *ago1-27* backgrounds. As expected, the ER marker (BiP) was enriched in the microsomal fraction, whereas the cytosolic enzyme UDP glucose pyrophosphorylase (UGPase) was absent, yet enriched in the soluble fraction (Fig. 6G). While in WT, FBW2- mediated AGO1 degradation was mainly visible in the soluble fraction, in the *ago1-27* background, the abundance of the mutated AGO1-27 protein decreased in both soluble and microsomal fractions after β-Es treatment. Note that FBW2 was also present in both fractions. From these results, we conclude that FBW2 associates with both soluble and membrane-bound AGO1, to trigger its degradation via a process that requires CDC48 activity.

### FBW2 degrades preferentially AGO1 when its loading is compromised

We noted that the degradation of AGO1 by FBW2 was more effective in transient expression assays than in stable transformed lines (Fig. 1; Fig. 2A; Fig. S3). A possible explanation of this phenomenon could be that most AGO1 is still unloaded when transiently expressed (Csorba et al., 2010), suggesting a preference of FBW2 for this form. To further address this question, we transiently co-expressed *AGO1* and *FBW2* with or without a construct carrying the inverse repeat of *GFFG*, which is known to produce functional siRNA (Himber et al., 2003). We reasoned that transient co-expression of the *GFFG* construct with AGO1 would foster its loading and could thus protect it from FBW2. However, AGO1 degradation by FBW2 was found only slightly attenuated in presence of *GFFG* (Fig. S5A-B). By contrast, when we co-expressed the VSR P19 that specifically binds 19-21-nucleotide double-stranded small RNA (Csorba et al., 2015) and could increase the unloaded pool of AGO1, the degradation of AGO1 was more efficient. To further support this observation, we also constitutively overexpressed the P19 protein in the Arabidopsis XVE:3HA-FBW2/*fbw2-1* mutant background to deplete at least a fraction of the pool of endogenous small RNA. In agreement with the transient expression assays, AGO1 degradation by FBW2 became far more effective when P19 was co-expressed than in the Col-0 background (compare Fig. S5C and Fig. S3A). Note also the enhanced AGO2 protein level upon FBW2-mediated AGO1 depletion.

Next, we took advantage of Arabidopsis mutants affecting the production or stability of small RNA to further investigate the possible impact of AGO1 loading on its degradability by FBW2. Therefore, we crossed *fbw2-4* and *FBW2OE* mutant lines with *hyl1-2* and *hen1-6* mutants. DRB1/HYL1 mediates the processing of most miRNA precursors (Kurihara et al., 2006), whereas the RNA methyltransferase HEN1 is critical for small RNA stability (Li et al., 2005) (Ren et al., 2014). Overexpression of *FBW2* in both mutant backgrounds revealed stronger growth and developmental defects than in the single mutants (Fig. 7A). At the molecular level, these phenotypes correlated well with decreased AGO1 protein levels (Fig. 7B) indicating that under conditions in which small RNA accumulation is affected, AGO1 becomes more prone to degradation by FBW2.

**Fig. 7.**
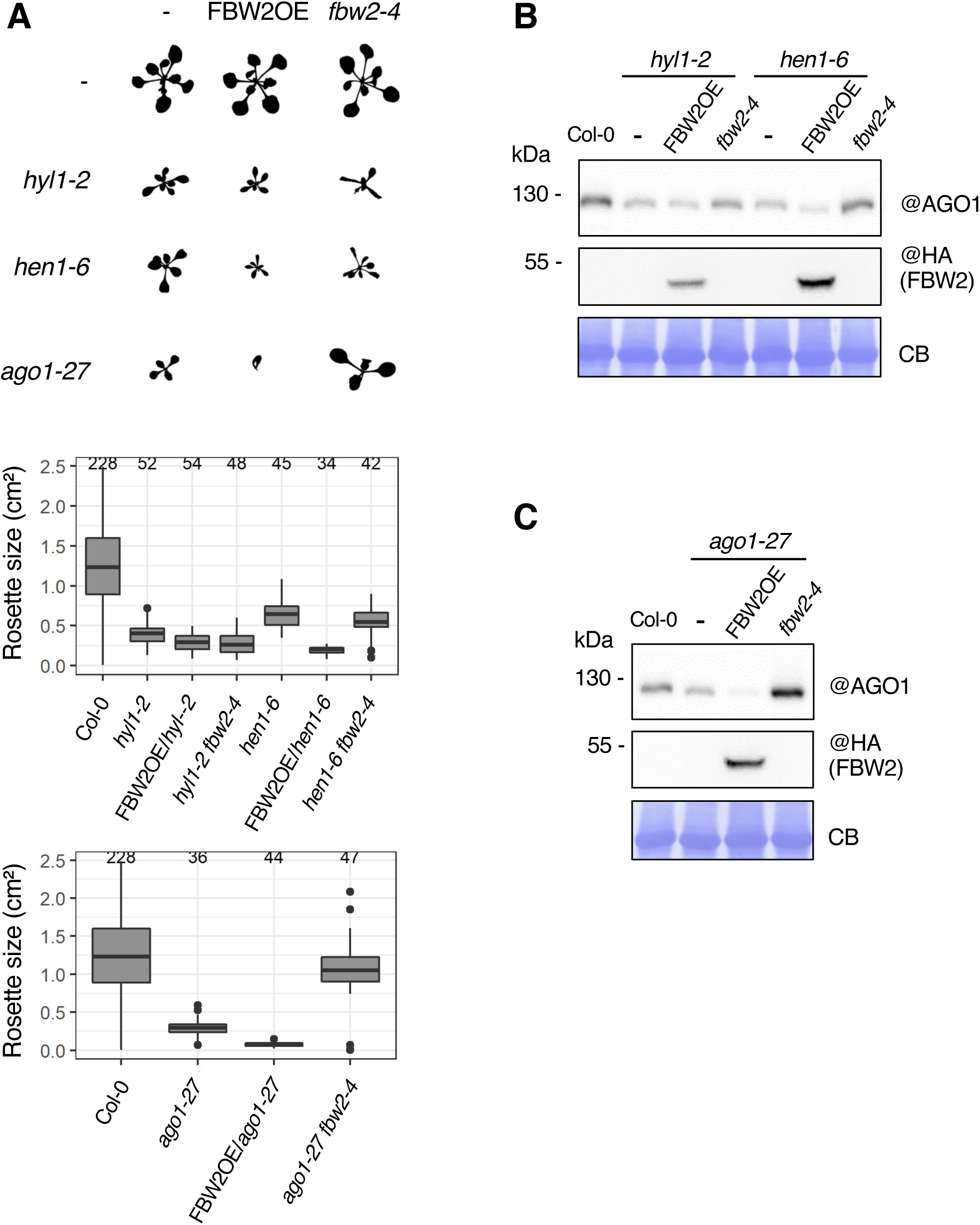
Effects of *FBW2* overexpression or loss-of-function in silencing mutants. (**A**) Upper panel: Shape imprints from 17 day-old seedlings of Col-0, *hyl1-2, hen1-6, ago1-27* and their crosses with *fbw2-4* or 35S:3HA-FBW2 (FBW2OE line 10) as indicated. Bottom panel: Measurements of the rosette area of the same seedlings represented as Tukey boxplots. The number of plants measured is indicated on top of the graphs. (**B-C**) Western blot of protein extracts from the same seedlings as indicated in (A). Coomassie blue (CB) staining was used as loading control. The “@” symbol indicates hybridization with the corresponding antibodies.

### Stabilized AGO1 in mutants impaired in small RNA accumulation is deleterious for plant development

In line with the previous report of (Earley et al., 2010), we observed that AGO1 protein level is at least partially restored in *hyl1-2 fbw2-4* and *hen1-6 fbw2-4* double mutants (Fig. 7B; Fig. S6A). Strikingly, despite this increased amount of AGO1 protein, we noticed that the growth and developmental phenotype of these double mutant plants were significantly exacerbated when compared to the single mutants, suggesting that the stabilized AGO1 protein became somehow toxic (Fig. 7A; Fig. S6B). This strongly contrasts with the situation of *ago1-27 fbw2-4* double mutant for which the increased AGO1-27 steady state protein level at least partially rescued the growth defect of the mutant (Fig. 7A, C). To better understand the reason of the apparent toxicity of stabilized AGO1 in *hyl1-2* and *hen1-6* mutant backgrounds, we first investigated the behaviour of the protein in the formation of protein complexes. For these experiments we chose to work with the *hyl1-2* mutant, as *hen1-6* was nearly sterile. It has been shown that AGO1-RISC complexes are present in high and low molecular weight complexes (Baumberger and Baulcombe, 2005)(Csorba et al., 2010), but only the low molecular weight complex exhibits the slicing activity, as in animals (Nykänen et al., 2001). We thus examined the molecular weight of AGO1-based RISCs in *hyl1-2 fbw2-4* seedlings by gel filtration, and the elution fractions were analysed by western blot (Fig. 8A; Fig. S7). As expected, Col-0 exhibited both high and low molecular weight AGO1-based RISCs. The *fbw2-4* single mutant behaved similarly to WT Col-0 showing both types of complexes. In contrast, the *hyl1-2* single mutant mainly presented low molecular weight RISCs, suggesting that the high molecular weight AGO1 complexes depend on miRNA accumulation. Interestingly, in the *hyl1-2 fbw2-4* double mutant, at least a fraction of the high molecular weight AGO1 complexes were re-established. Accordingly, miR159 co-fractionated with both the low and high molecular weight AGO1-containing complexes in Col-0, while in *hyl1-2* and *hyl1-2 fbw2-4* the miR159 was barely detected in any fraction probably because of its impaired synthesis (Fig. 8B). This observation does not hold true for a microRNA like miR168, whose abundance is only marginally affected by the loss of *HYL1* (Szarzynska et al., 2009). Based on these observations, we hypothesized that when miRNA availability is compromised and AGO1 degradation is impaired, as in the *hyl1-2 fbw2-4* double mutant, unconventional AGO1-bound RNA may be incorporated in RISCs, as supported by our gel filtration assay, and may ultimately become problematic for the plant, as indicated by the more severe phenotype in *hyl1-2 fbw2-4* and *hen1-6 fbw2-4* double mutants (Fig. 7A; Fig. S6).

**Fig. 8.**
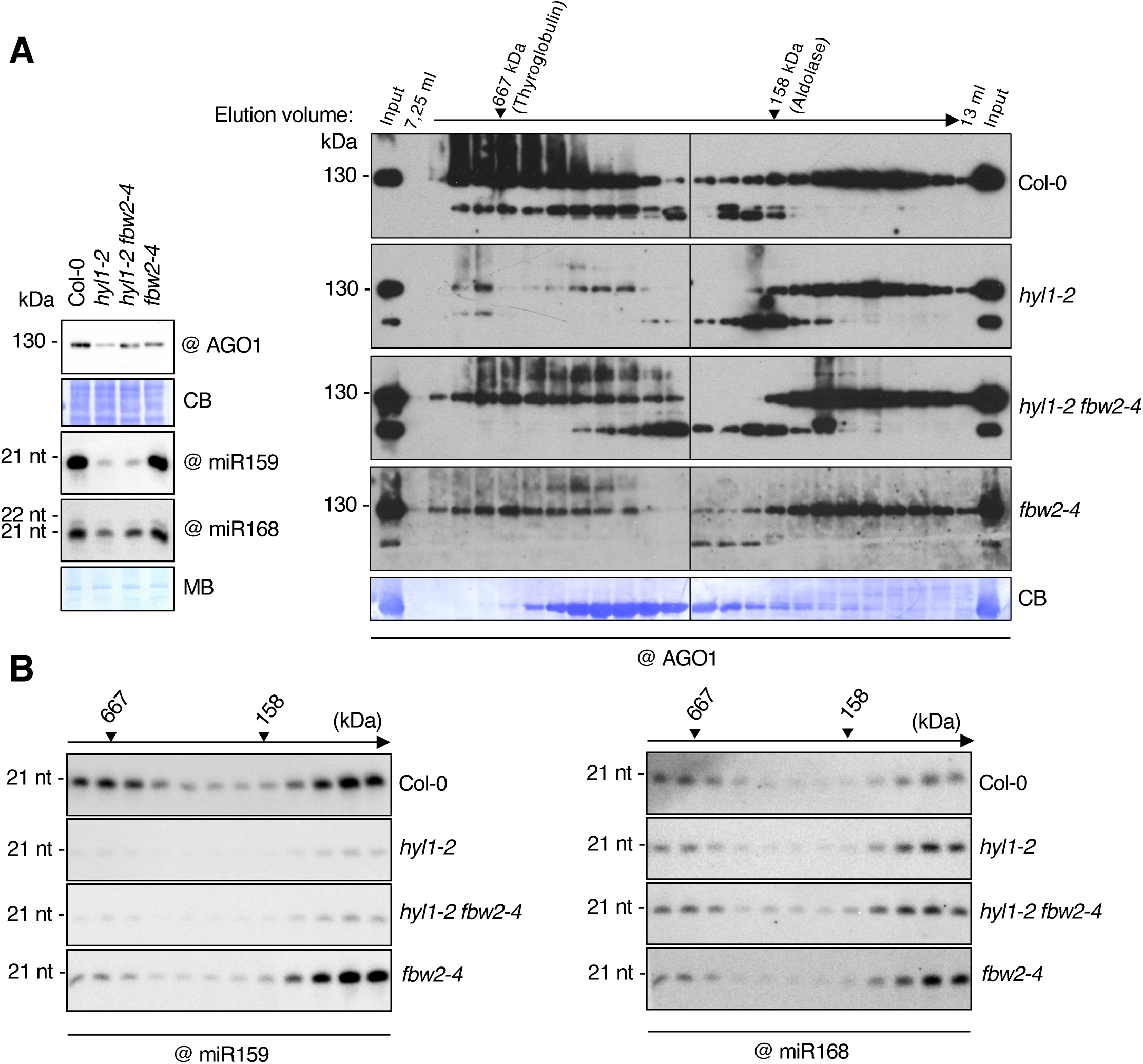
Loss of *FBW2* restores high molecular weight AGO1 complexes in *hyl1-2*. **(A)** Gel filtration analysis of AGO1-based RISC complexes in Col-0, *hyl1-2*, *hy1-2 fbw2-4* and *fbw2-4* 13 day-old seedlings. Proteins of known molecular weight are shown on top of the blot. Coomassie blue (CB) staining was used as loading control and “@” indicates hybridization with the AGO1 antibody. On the left panel are shown protein and small RNA analysis of the input fraction prior to gel filtration. Methylene Blue (MB) staining of the membrane was used as loading control. (B) Small RNA analysis from even fractions, spanning the same range. For this analysis, 10μg of the RNA per lane was loaded. The “@” symbol indicates hybridization with the indicated oligonucleotide probes.

To better characterize the global RNA-binding activity of AGO1 in *hyl1-2* versus *hyl1-2 fbw2-4*, we immunoprecipitated AGO1 from the different genetic backgrounds and indiscriminately labelled the incorporated RNA by replacing their 5’ phosphate with a radioactive one, using Polynucleotide Kinase (PNK) (Fig. 9A-B). As expected, the *hyl1-2* mutant showed a reduced amount of miRNA loaded in AGO1 while the pattern of RNA associated to AGO1 in *fbw2-4* was similar to Col-0. Remarkably, the amount of 21/22 nt long small RNA bound to AGO1 is re-established in the *hyl1-2 fbw2-4* double mutant and, in addition, AGO1 became more loaded with 24 nt long small RNA species in this genetic background.

**Fig. 9.**
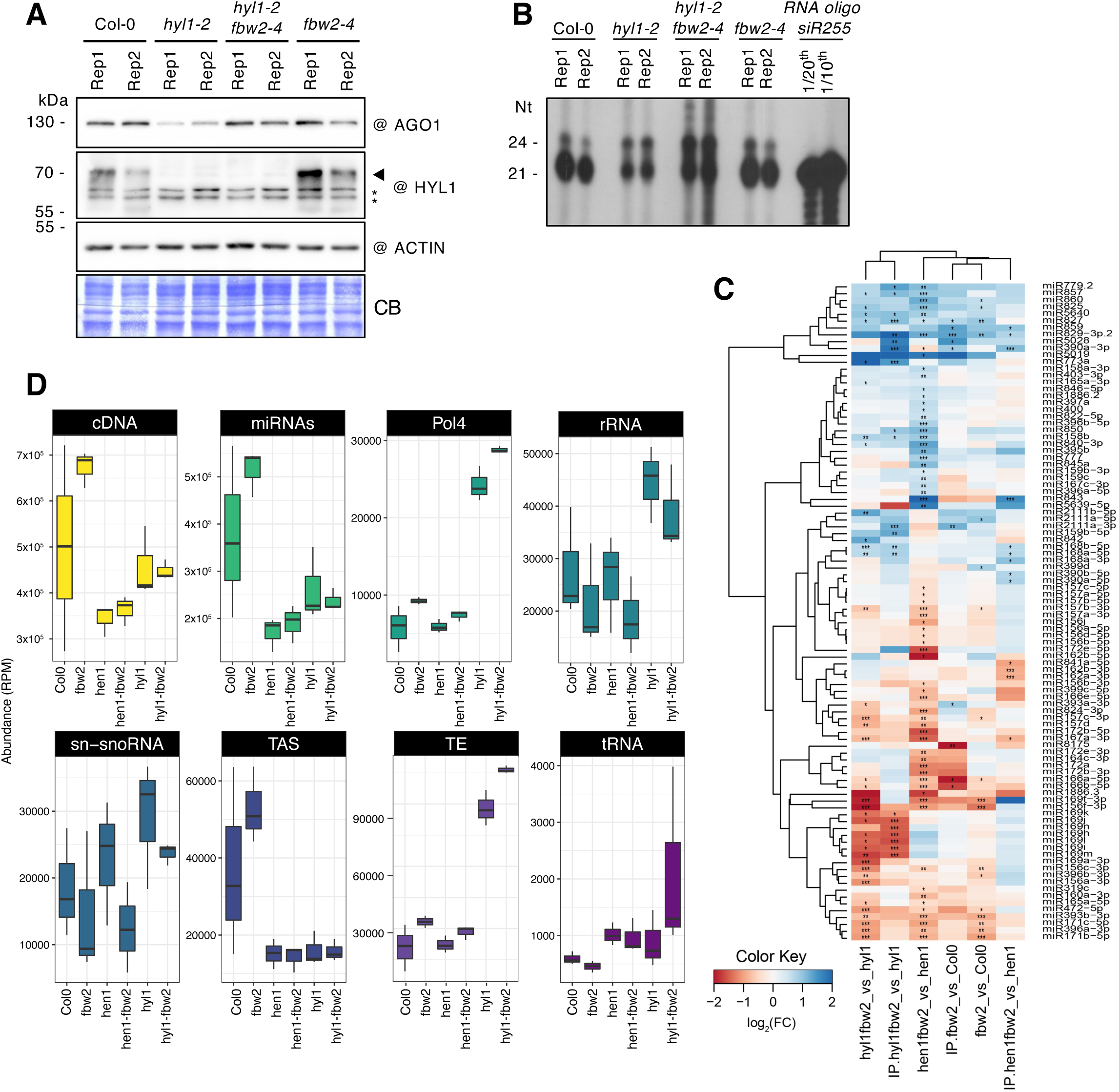
Loss of *FBW2* modifies AGO1 loading in *hyl1-2*. **(A)** Western blot of total protein extracts from 2 week-old seedlings from Col-0, *hyl1-2, hy1-2 fbw2-4* and *fbw2-4* mutants. Two biological replicates (1 and 2) were made. Coomassie blue (CB) staining and ACTIN were used as loading controls and “@” indicates hybridization with the corresponding antibodies. The arrow indicates the HYL1 protein band while the * symbol indicates aspecific cross-reacting bands. (B) Denaturing polyacrylamide gel of small RNA from immunoprecipitated AGO1 from the same protein extracts shown in (A). RNAs were indiscriminately labelled by replacing their 5’ phosphate with a radioactive one using Polynucleotide Kinase (PNK). An oligo corresponding to the siR255 serves as control for small RNA size. (C) Relative abundance of miRNA in AGO1 IP samples with significant differential expression in single and double mutants compared to wild-type (Col-0) and single mutants; the relative abundance is expressed as a heat map (see Key at the bottom), with the samples being compared indicated below each heat map (*Q value # 0.05, **Q value # 0.01 and **Q value #0.001). (D) Boxplot representing the abundance of reads (in reads per million) mapping to eight different features of the Arabidopsis genome TAIR 10. These include the following from left to right: cDNA; mature miRNA; siRNA precursors dependent on Pol4; ribosomal RNAs (rRNAs); small nuclear and small nucleolar RNA (snRNA and snoRNA); TAS precursors; Transposable elements (TEs); tRNA-derived small RNA (tRNA); RPM, reads per million.

### Identification of small RNA loaded into stabilized AGO1 and their targets

To get more insights into the identity of small RNA in the context of the *fbw2-4* mutation, we performed deep-sequencing analyses on total small RNA and AGO1-associated small RNA in Col-0 and five different mutant backgrounds (e.g., *fbw2-4, hyl1-2* and *hen1-6* single, and *hyl1-2 fbw2-4* and *hen1-6 fbw2-4* double mutants) (Table S2). AGO1 protein levels in the different mutants were verified beforehand by western blot (Fig. S8). As expected, AGO1 protein amounts were lower in the three replicates of *hyl1-2* and *hen1-6*, but at least partially re-established in the double mutants when combined with *fbw2-4*.

As previously described, we observed that the size distribution of total small RNA is significantly altered in *hen1* and *hyl1* mutant backgrounds (Yu, 2005)(Zhai et al., 2013)(Li et al., 2005)(Kurihara et al., 2006) (Fig. S9A). However, we noted that the mutation of *fbw2-4* did not alter the size distribution in any of the three studied backgrounds, Col-0, *hen1-6* or *hyl1-2*. The two predominant peaks at 21- and 24-nt observed in the WT and *fbw2-4* mutants completely disappeared in the *hen1-6* and *hen1-6 fbw2-4* mutants, while the 21-nt peak disappeared and the 24-nt peak increased in the *hyl1-2* and *hyl1-2 fbw2-4* mutants. Regarding the AGO1-associated small RNA, we observed a similar situation, with a significant decrease of 21-mers in *hen1-6* and *hen1-6 fbw2-4* mutants, and a significant increase in 24-mers in *hyl1-2* and *hyl1-2 fbw2-4* mutants (Fig. S9B), as observed for AGO1-associated small RNA labelling (Fig. 9A-B). Next, we analysed the miRNA differential accumulation (DA) in the single and double mutants compared to WT, and in the double mutants compared to single mutants (Fig. 9C; Fig. S10). Very few miRNA were DA when comparing the *fbw2-4* mutants with the WT; only 18 miRNA in total RNA samples (six up- and 12 down-accumulated) and 11 in the AGO1 IP samples (eight up- and three down-accumulated) (Fig. 9C). We also found few DA miRNA when comparing the double mutant *hyl1-2 fbw2-4* to *hyl1-2* and *hen1-6 fbw2-4* to *hen1-6*, in both total RNAs and AGO1 IP samples (Fig. 9C; Fig. S10).

To better understand the phenotype observed for our mutants, we evaluated the genomic origin of all small RNA reads in each mutant, and we focused on the new category of 24-mers bound to AGO1. We mapped all the reads from each mutant to different features in the genome. As expected, we observed that the *hyl1-2 fbw2-4* and *hyl1-2* mutants accumulated lower levels of reads derived from miRNA and *trans*-acting siRNA (*TAS*) genes (Fig. S11). However, these mutants also accumulated a higher number of small RNA originating from Pol IV products and transposable elements (TEs), and, when focusing on the 24-nt long small RNA, these were loaded into AGO1 (Fig. 9D; Fig. S9B). Furthermore, these mutants had more rRNA-derived 24-nt small RNA loaded in AGO1 compared to WT and the other mutants (Fig. 9D). We also generated Parallel Analysis of RNA End (PARE) libraries from the same material as for the small RNAseq libraries (Table S2), allowing us to identify 4301 small RNAs/targets. We only considered small RNA/target signatures present in all the biological replicates, and produced by small RNA identified in AGO1 IP libraries between 19 and 24 nt in length. We observed that, as expected, most of the small RNA/target signatures were shared between all the samples, and were a product of miRNAs and tasiRNA (Fig. 10A). In our conditions, tRF and snRNA and snoRNA did not seem to play a major role in producing small RNA with a target. We also observed a subset of specific signatures for each genotype, corresponding mostly to targets that were not present in the other genotypes. To understand how the presence or absence of these “new” or genotype-specific targets could impact the plant, we analysed the gene ontology (GO) term enrichment in each of the tested genotypes. In our analysis, we considered only GO terms that are statistically significantly enriched when comparing the mutants to the WT (Fig. 10B). We observed that the double mutants *hyl1-2 fbw2-4*, compared to Col-0 or compared to the single mutants *hyl1-2*, has an enrichment in targets belonging to GO categories such as response to stress, biosynthesis of organic substances and cellular metabolic and biosynthetic processes, as well as a depletion of genes belonging to response to radiation and light stimulus. These results highlight clear differences in small RNA/target signatures in the double versus single mutants, which could at least in part explain their phenotype.

**Fig. 10.**
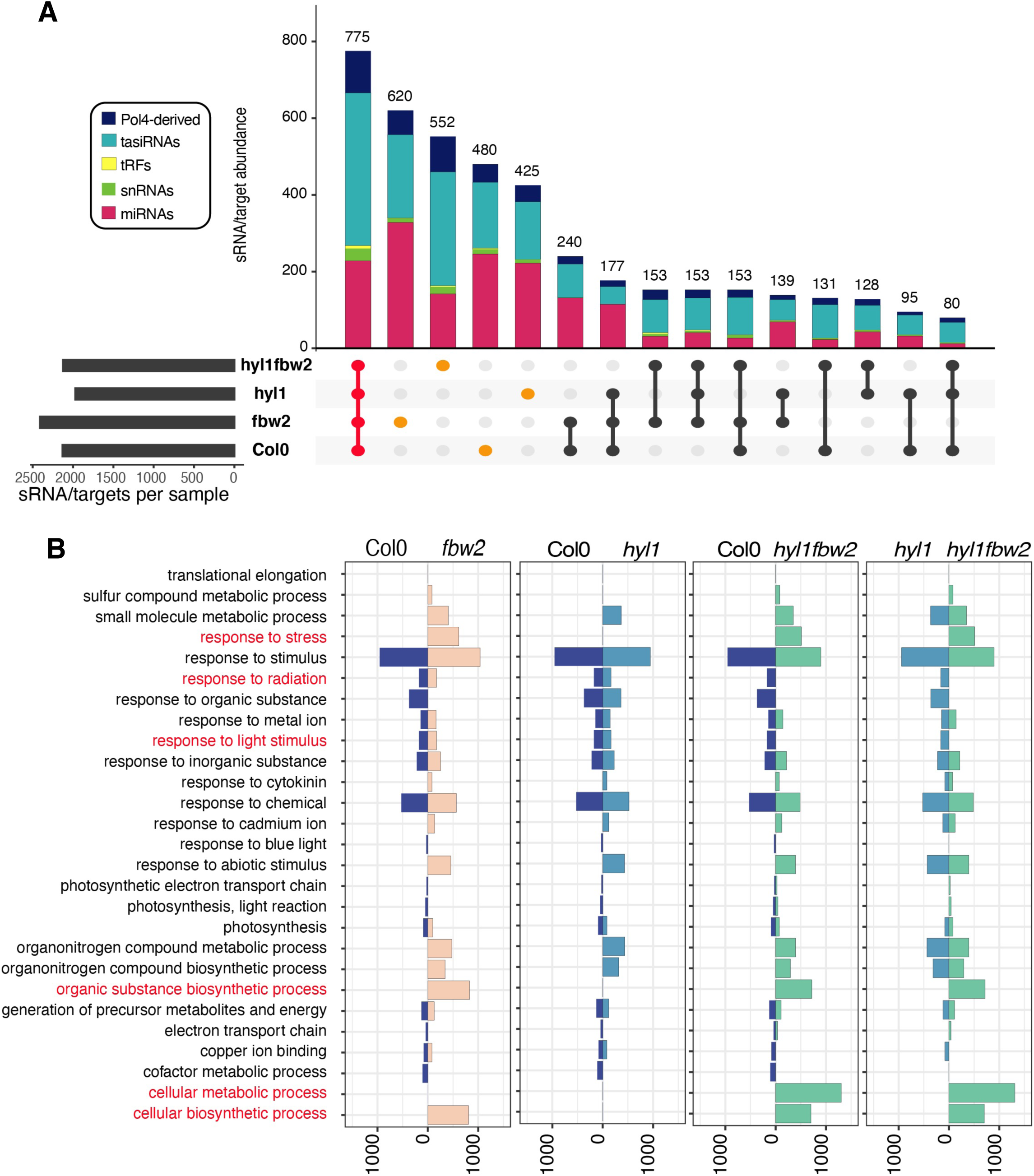
Loss of *FBW2* modifies AGO1 small RNA-mediated targeting. **(A)** Visualization of the intersection of small RNA/target signatures among the different mutants, represented with an UpSet plot. The upper panel shows a vertical bar plot with the number of signatures included in each intersection, colour coded into five small RNA categories (including from top to bottom: siRNA precursors dependent on Pol4; siRNA from TAS precursors (tasiRNA); tRNA-derived sRNA (tRF); small nuclear and small nucleolar RNA (snRNA); miRNA). The bottom left panel shows a horizontal bar chart with the number of signatures included in each set. The bottom right panel indicates which intersections and their aggregates are being considered in each case. Labelled in red are the signatures common to all samples, and in orange are the signatures unique to each mutant. (B) Vertical bar plot representing the number of genes (horizontal scale) included in each gene ontology (GO) term (vertical scale). Only significantly enriched GO terms are represented for each of the tested genotypes. From left to right we represent Col-0 versus *fbw2* mutants, Col-0 versus *hyl1* mutants, Col-0 versus *hyl1bfw2* double mutants, and *hyl1* mutants versus *hyl1 fbw2* double mutants.

## Discussion

### Similarity and divergence of viral P0 and endogenous FBW2 F-box proteins

While the function of AGO proteins and their bound small RNA have been extensively studied in various biological processes across several organisms (Meister, 2013), their regulation at the post-translational level is less understood. Our laboratory and others have previously unravelled the mode of action of a viral encoded F-box protein P0 from poleroviruses, which promotes the degradation of AGO1 and thus presumably impairs RNA-based anti-viral immunity (Baumberger et al., 2007)(Bortolamiol et al., 2007) (Csorba et al., 2010). Because viruses usually hijack host cell machineries, it is conceivable that P0 could usurp the function of an endogenous F-box protein such as FBW2 during infection. However while both F-box proteins target AGO proteins, our data indicate clear differences in their mode of action, specificity and regulation.

First, both F-box proteins do not recognize their substrates through the same structural domains. Thus, the degron motif recognized by P0, which has been precisely mapped, is localized in the conserved DUF1785 domain of AGO1 and this degron can also confer P0-mediated degradation to other AGO proteins (Derrien et al., 2018). Although the degron necessary for FBW2-mediated AGO1 degradation has not yet been defined precisely, it is located in the MID-PIWI domain. Second, P0 does not only mediate AGO1 turnover, but also triggers the degradation of at least AGO2 and AGO4 when transiently expressed in tobacco leaves (Baumberger et al., 2007) and in stable transformed Arabidopsis lines (Derrien et al., 2018)(Trolet et al., 2019). This broad activity of P0 on several AGOs is likely key for its activity as a VSR, as besides AGO1, at least AGO2, AGO5 and AGO7 possess antiviral activities against RNA viruses (Qu et al., 2008) (Takeda et al., 2008). On the contrary, FBW2 acts specifically on AGO1 and possibly other members of its clade. Third, both F-box proteins are also regulated differently at the post-translational level. P0 interacts with AGO1 on the ER and both proteins are codelivered to the vacuole where they are degraded via an autophagy-like process (Michaeli et al., 2019). This is clearly not the case for FBW2, which is degraded by the proteasome. Since MLN4924, a drug that inhibits CUL1 neddylation, also blocks its degradation, it is possible that FBW2 catalyses its own ubiquitylation, as has been reported for other F-box proteins (Zhou and Howley, 1998) (Galan and Peter, 1999).

Finally, P0 proteins from different poleroviruses are unstructured proteins, though at least some carry GW/WG-like motifs that are required for suppression of RNA silencing (Zhuo et al., 2014). Interestingly, a structural feature in the C-terminal domain of FBW2 is the enrichment in tryptophan-residues showing some characteristics of GW-repeat proteins. Although only one GW motif could be predicted using bioinformatic tools, other W and neighbouring residues, as well as their spacing, suggest the presence of such a domain in FBW2. This motif, also defined as an AGO hook, was first identified in the mammalian GW182 protein and shown to interact with AGO proteins and be essential for miRNA-guided gene silencing in different animal organisms (reviewed in (Pfaff and Meister, 2013)). In Arabidopsis, the protein SUO involved in miRNA-mediated translational repression has been proposed to act as a functional analogue of GW182 (Yang et al., 2012). Nevertheless, whether the GW-repeats of SUO are required for the interaction with AGO1 is presently unknown. Note that other GW-repeat containing proteins have been predicted in Arabidopsis (Karlowski et al., 2010) and one of them, SDE3, which is involved in different silencing pathways, interacts preferentially with AGO1 through its GW motifs (Garcia et al., 2012). For FBW2, the mutation of at least four W residues significantly impaired the capacity of the protein to degrade AGO1. As FBW2 recognizes the MID-PIWI domain for degradation and since these domains of human Ago2 are known to be important for GW protein binding and anchoring the 5’ end of the guide RNA (Schirle and MacRae, 2012), it will be interesting to further investigate the structural and functional role of the FBW2 C-terminal domain.

### Mechanism of FBW2-mediated AGO1 degradation

The stability of AGO proteins has been extensively linked to their loading state. For instance, it has been shown that the inhibition of HSP90 activity, which is required for AGO loading across eukaryotes (Iwasaki et al., 2010)(Iki et al., 2010), destabilizes human Ago1 and Ago2 proteins (Johnston et al., 2010). Accordingly, mutations affecting small RNA availability also destabilize AGO proteins in Arabidopsis (AGO1), Drosophila (Ago1) and mammals (Ago2) (Derrien et al., 2012)(Smibert et al., 2013)(Martinez and Gregory, 2013). The fact that mutation of *FBW2* restores AGO1 protein levels in various mutants affecting small RNA biogenesis, accumulation or loading of AGO1 ((Earley et al., 2010) and our work) strongly argues that this F-box protein targets the unloaded form of AGO1. This is further supported by the observation that the degradation of AGO1 by FBW2 was more effective in transient expression assays, a situation in which most AGO1 is still unloaded (Csorba et al., 2010), while in stable transformed Arabidopsis lines, the presumably small RNA loaded AGO1 was more resistant to this degradation. Note that in transient expression assays, the co-expression of a *GFFG* construct generating siRNAs and thus fostering AGO1 loading was unable to block its degradation by FBW2, but only attenuated it. Nevertheless, once we constitutively co-expressed P19 in Arabidopsis, which binds to both siRNA and miRNA/miRNA* duplexes, AGO1 became more susceptible to FBW2-mediated degradation. This situation is reminiscent to Drosophila, where unloaded Ago1 can be rescued from degradation by synthetic miRNAs but not siRNAs (Smibert et al., 2013). Interestingly it has recently been shown that the unloaded form of Drosophila Ago1 is recognized and subsequently degraded by a RING-type E3 ubiquitin ligase named Iruka (Kobayashi et al., 2019a). This work showed that Iruka ubiquitylates Lys514 in the L2 linker of Ago1, which is only accessible in its empty state. However, other E3 ubiquitin ligases are able to target loaded forms of AGO proteins. Hence, it was recently found that ZSWIM8, a cullin3-RING ubiquitin ligase (CRL3) adaptor protein, ubiquitylates human Ago2 when engaged with a TDMD (target-directed microRNA degradation) target, leading potentially to proteasomal degradation of the miRNA-containing complex (Han et al., 2020)(Shi et al., 2020).

Notably, FBW2 may potentially also recognize the loaded form of AGO1. For instance, we observed that FBW2 associates not only with soluble but also membrane-bound AGO1; this form of AGO1 is likely loaded and bound to its target RNA. Moreover, it was recently proposed that FBW2 may target AGO1 when it is bound to an mRNA under prolonged conditions, such as in the presence of non-cleavable artificial miRNA target mimics, expected to prolong the time of AGO1-target interaction (Ré et al., 2019). Though the direct involvement of FBW2 in this mechanism still remains to be demonstrated, it will be interesting to determine whether AGO1 would undergo conformational changes on its target RNA allowing its recognition by the F-box protein. It is also tempting to speculate that certain mutations, such as in *ago1-27*, leading to an increased degradation rate of the AGO1-27 protein by FBW2, would be subjected to a similar regulation.

A question not yet fully solved is by which proteolytic mechanism does FBW2 mediate AGO1 degradation. In animal cells, AGOs have been shown to undergo proteasomal degradation (Johnston et al., 2010)(Chinen and Lei, 2017)(Han et al., 2020)(Shi et al., 2020), but also degradation via lysosomes or even selective autophagy (Gibbings et al., 2012)(Martinez and Gregory, 2013). Interestingly, at least in Drosophila S2 cells, recent work elucidated a novel pathway in which valosin-containing protein (VCP/p97/Cdc48) directs degradation of ubiquitylated empty Ago1 through the Ufd1-Npl4 adaptor subunits (Kobayashi et al., 2019b). In Arabidopsis, our pharmacological approach revealed that the proteasome is not the main route for FBW2-mediated AGO1 decay, though a small fraction of AGO1 undergoes FBW2-dependant proteasomal degradation as supported by a slight stabilisation of AGO1 protein and the appearance of a cleavage product in the presence of Bortezomib, as well as a significant enrichment of proteasomal subunits in the FBW2 IP. Similarly, by using the *atg7-2* mutant resulting in an inability to deliver autophagosomes to the vacuole (Marshall and Vierstra, 2018), bulk autophagy was also excluded as the prime mechanism for FBW2- mediated AGO1 degradation. The fact that the drug MLN4924, which blocks the activity of CULLIN-based E3 ubiquitin ligases, had a more pronounced effect in blocking AGO1 degradation highlights the importance of ubiquitylation in this process. Since both E64d and CB-5083 significantly prevented AGO1 decay, we assume that FBW2 promotes AGO1 vacuolar degradation, via a pathway involving CDC48, which could be similar to the mechanism described in flies (Kobayashi et al., 2019b). Elucidating the molecular and cellular determinants of this pathway will represent an important goal in the future.

### Why is it important to degrade AGO1?

A key question to address is the physiological importance of AGO protein turnover by ubiquitin E3 ligases. In the fission yeast *Schizosaccharomyces pombe*, it was hypothesised that the accumulation of unloaded Ago1, which is involved in transcriptional silencing, might be problematic to cells as it could generate inactive complexes poisoning the activity of small RNA–programmed RNA-induced transcriptional silencing (RITS) complexes (Holoch and Moazed, 2015). As discussed above, in Drosophila, the ubiquitin E3 ligase Iruka eliminates the empty form of Ago1 (Kobayashi et al., 2019a). In this work, the authors suggested that this mechanism would be particularly relevant for dysfunctional forms of Ago1, potentially originated from translational errors or incorrect folding and locked in an empty state. In the future, *Iruka* loss-of function flies might reveal the *in vivo* importance of such a protein quality control mechanism.

In Arabidopsis, *FBW2* loss of function does not affect plant growth or development under standard growth conditions nor the accumulation level of specific miRNA. In *fbw2* mutant plants, the AGO1 protein level was found only slightly increased, though this already significantly affected S-PTGS activity. Thus, one evident function of FBW2 is to work conjointly with the miR168 feedback loop (Mallory and Vaucheret, 2010) to maintain AGO1 homeostasis. Note, that the latter appears more decisive as the expression of miR168-resistant *AGO1* mRNA induces severe development defects in Arabidopsis (Vaucheret et al., 2004). Another possible function for FBW2 could be its involvement in the degradation of mutated and dysfunctional forms of AGO1, as proposed for Drosophila (Kobayashi et al., 2019a). This is supported by the observation that the AGO1-27 mutant protein was more susceptible to FBW2-mediated degradation, if compared to native AGO1.

Interestingly, AGO1 post-translational control by FBW2 revealed its importance particularly under certain conditions. Indeed, when the *fbw2* mutation was combined with *hyl1-2* or *hen1-6* mutants, respectively affecting the production and stability of small RNA (Kurihara et al., 2006) (Li et al., 2005) (Ren et al., 2014), the stabilised AGO1 protein enhanced the growth and developmental phenotype of the single mutants. Deep-sequencing analyses of small RNA in the different mutant background revealed that under these conditions, AGO1 associates *in vivo* with small RNA, derived from categories yielding few small RNA in a WT context. Strikingly, the loading of some illegitimate small RNA in stabilised AGO1 leads to the cleavage of new target genes belonging to diverse pathways including stress responses and also cellular metabolic processes. This abnormal targeting likely contributes to the enhanced phenotype observed in the *hyl-1 fbw2* double mutant. Whether the control of AGO homeostasis by E3 ubiquitin ligases to avoid off-target cleavage also operates in other organisms, such as mammals, or is unique to plants, will need further investigation.

## Materials and Methods

### Plant material, growth conditions and treatments with chemicals

*Arabidopsis thaliana* ecotype Colombia as well as *Nicotiana benthamiana* (for transient expression) were used in this study. The following Arabidopsis mutants, *ago1-27* (Morel et al., 2002), *ago1-57* (Derrien et al., 2018), *fbw2-1* (Earley et al., 2010), *fbw2-4* (SALK_144548), *atg7-2* (GK-655B06); *hen1-6* (SALK_090960), *hyl1-2* (SALK_064863), *sqn-1* (Smith et al., 2009) and *sgs2-1* (Mourrain et al., 2000) were used. The XVE:P0-myc, Hc1, L1 and Hc1*/sqn-1* and Hc1*/4mAGO1* stable lines have been described previously (Derrien et al., 2012)(Elmayan et al., 1998)(Martínez De Alba et al., 2011). T-DNA plant transformation was performed using the floral dip method (Clough and Bent, 1998).

For *in vitro* culture conditions, Arabidopsis seeds were surface-sterilized using ethanol and plated on MS agar (MES-buffered MS salts medium [Duchefa, Murashige & Skoog medium inc. vitamins/MES- MO255)], 1% sucrose, and 0.8% agar, pH 5.7). The seeds were then stratified for 2 days at 4°C in the dark and then transferred in 16h-light/8h-dark (20,5/17°C, 70% humidity) growth chamber, under fluorescent light (Osram Biolux 58W/965). Unless otherwise specified, seedlings were transferred in liquid MS (MES-buffered MS salts medium [Duchefa, MO255], 1% sucrose, pH 5.7) and acclimated for 24 hours prior to chemical treatments.

For P0-myc and XVE:3HA-FBW2 induction during plant growth, MS-agar plates were supplemented with 10μM β-estradiol, while for mock treatment, an equal amount of DMSO was used. Plates were then handled as indicated above, and seedlings were harvested at 7 to 8 days or as indicated after sowing for protein content analysis or 9 to 10 days after sowing for aerial and root growth measurements.

For kinetic induction of XVE:3HA-FBW2, seedlings were grown as indicated above for 8 to 12 days, then transferred into liquid MS medium (Duchefa, MO255) +1% sucrose in sterile conditions. Liquid MS medium was then replaced with either MS + DMSO (mock) or MS + 10μM β-estradiol. Similarly, for drug treatment, the different Arabidopsis lines (specified in the figures) were grown for 8 to 12 days, then transferred into liquid MS medium (Duchefa, MO255) +1% sucrose in sterile conditions. Liquid MS medium was then replaced with either MS + DMSO (mock=M), MS + 20μM MLN4924, or MS +100 μM Bortezomib (Bortz), or MS + 50 or 100μM E64d, or MS + 20μM CB-5083 (CB). In the specific case of XVE:3HA-FBW2 lines, 10μM β-estradiol was supplemented in the liquid medium in addition to the different drugs. Immerged seedlings were then left in the growth chamber under slow agitation for the indicated period of time before harvest.

For IP-MS experiments, Col-0, FBW2OE and FBW2O/*ago1.27* Arabidopsis lines were grown for 8 days on MS-agar plates then transferred into liquid MS medium + 20μM MLN4924 for 20 hours before harvesting.

### Analysis of leaf area and lateral roots

For leaf area analysis, seedlings were grown in soil. The seeds were stratified in water and darkness for 48 hours. Plastic pots (7×7×7cm) were filled with 62g of soil (Hawita Fruhstorfer erde) and watered with tap water to reach a relative humidity of 2.2g_H20/gsoil_ (RWC 69%). Three seeds were sown in the middle of the pot. The pots were covered with a plastic foil to maintain the humidity level and placed in a 16h-light (21°C) and 8h-darkness (18°C) regime. At 4 days after stratification (DAS), the plastic film was removed, and at 5 DAS plants were thinned out to keep one plant per pot. The pots were watered every 2 to 3 days and maintained at a relative soil humidity of 2.2_gH20/gsoil_. Plant size at 22 DAS was determined by dissecting every leaf and placing it from oldest to youngest on a petri dish with 1% agar. The plates were photographed and the leaf area was measured with ImageJ v1.45 (NIH; https://rsb.info.nih.gov/ij/). Ten plants per line were used.

For lateral root measurements, seedlings were grown on vertical petri dishes with MS medium as described above. After eight days of growth, the number of lateral roots of each seedling was counted under the binocular. Only macroscopically visible lateral roots were counted (stage >VIII). At least ten seedlings per line were analysed.

### Plasmid constructions

The 35S:P19, 35S:GUS, and 35S:P0-myc, 35S:Flag-AGO1, 35S:Flag-AGO2, 35S:Flag-AGO3, 35S:Flag-AGO4, 35S:Flag-AGO5 constructs have been described in (Baumberger et al., 2007). The XVE:P0-6myc, p35S-GFP, p35S-GFFG constructs have been described in (Bortolamiol et al., 2007). The 35S:CFP-AGO1 construct was described in (Derrien et al., 2018).

The AGO1 WT constructs (pENTRY(Zeo)-AGO1) was generated by PCR amplification from the Arabidopsis AGO1 cDNA with the oligonucleotide primers listed in Table S3. Amplicons containing the attB sites were recombined into pDONR Zeo plasmids (Invitrogen). They were then transferred into the binary vector pK7WGF2, pB7WGC2 and pH7WGF2 (Karimi et al., 2005) by Gateway LR reaction to create the final N-terminal GFP, CFP or RFP fusion placed under the regulation of the 35S promoter

The AGO1 deletion constructs (pENTRY(221)-polyQ-ND, pENTRY(221)-ND-DUF, pENTRY(221)-ND-PAZ, pENTRY(221)-DUF, pENTRY(221)-DUF-PAZ, pENTRY(221)- PAZ, pENTRY(221)-L2, pENTRY(221)-L2-MID, pENTRY(221)-MID, pENTRY(221)-MID-PIWI, pENTRY(221)-PIWI) were generated by PCR amplification from the Arabidopsis AGO1 cDNA with the oligonucleotide primers listed in Table S3. Amplicons containing the attB sites were recombined into pDONR 221 plasmids (Invitrogen). They were then transferred into the binary vector pK7FWG2 (Karimi et al., 2005) by Gateway LR reaction to create the final C-terminal GFP fusion placed under the regulation of the 35S promoter (pK7FWG2-polyQ-ND, pK7FWG2-ND-DUF, pK7FWG2-ND-PAZ, pK7FWG2-DUF, pK7FWG2-DUF-PAZ, pK7FWG2-PAZ, pK7FWG2-L2, pK7FWG2-L2-MID, pK7FWG2-MID, pK7FWG2-MID-PIWI, pK7FWG2-PIWI).

The Flag-NtAGO1 construct (pENTRY(221)-FlagNtAGO1) was amplified using primers listed in Table S3. The Flag-NtAGO1 sequence containing the attB sites was recombined into pDONR 221 plasmids (Invitrogen) and then transferred into the binary vector pB2GW7 (Karimi et al., 2005) by Gateway LR reaction to create the final NtAGO1 N-terminal Flag fusion placed under the regulation of the 35S promoter.

The 3HA-FBW2 construct was generated by PCR amplification from the FBW2 cDNA with the oligonucleotide primers listed in Table S3. Amplicons were cloned by restriction (BamHI-NotI) into pE2N plasmid (Dubin et al., 2008). Then, 3HA-FBW2 was transferred from pE2N-3HA-FBW2 into the binary vector pB2GW7 (Karimi et al., 2005) by Gateway LR reaction to create the final N-terminal 3HA fusion placed under the regulation of the 35S promoter.

The 35S:FBW2 construct was generated by PCR amplification from the FBW2 cDNA with the oligonucleotide primers listed in Table S3 and used for gateway recombination using respectively the pDONR-Zeo plasmid (Invitrogen). Then, FBW2 was transferred from pENTRY(Zeo)-FBW2 into the binary vector pB2GW7 (Karimi et al., 2005) by Gateway LR reaction to create the final FBW2 placed under the regulation of the 35S promoter (p35S: FBW2).

For microscopy analysis, we generated the pFBW2:Venus-FBW2 construct. The *FBW2* promoter, the *Venus* and *FBW2* coding sequences were amplified by PCR using primers listed in Table S3 and used for gateway recombination using respectively the pDONR-P4P1R, pDONR-221 and pDONR–P2RP3 plasmids (Invitrogen). Then, the pENTRY obtained: pEN-L4-PromFBW2-R1, pEN-L1-VENUS-L2 and pEN-R2-FBW2-L3 were transferred into the binary vector pH7m34GW (Karimi et al., 2005) by Gateway LR reaction to create the final FBW2 N-terminal Venus fusion placed under the regulation of the *FBW2* promoter (pFBW2:Venus-FBW2).

The FBW2 WT, deletion and mutation constructs (pENTRY(221)-FBW2, pENTRY(221)-delF-box (FBW2), pENTRY(221)-delCter (FBW2), pENTRY(221)- FBW2(mutF-box), pENTRY(221)-FBW2(W225A), pENTRY(221)-FBW2(W258A), pENTRY(221)-FBW2(W275A), pENTRY(221)-FBW2(W295A), pENTRY(221)- FBW2(W313A), pENTRY(221)-FBW2(W258A, W275A), pENTRY(221)- FBW2(W258A,W275A, W295A), pENTRY(221)-FBW2(W225A, W258A, W275A, W295A), pENTRY(221)-FBW2(W225A, W258A, W275A, W295A, W313A) were generated by PCR amplification from the *FBW2* cDNA with the oligonucleotide primers listed in Table S3. Amplicons containing the attB sites were recombined into pDONR 221 plasmids (Invitrogen). They were then transferred into the binary vector pGWB415 (Nakagawa et al., 2009) by Gateway LR reaction to create the final N-terminal 3HA fusion placed under the regulation of the 35S promoter.

The FBW2 genomic construct (pENTRY(221)-iFBW2) was amplified from genomic DNA by PCR amplification using primers listed Table S3. The *FBW2* genomic sequence containing the attB sites was recombined into pDONR 221 plasmids (Invitrogen) and then transferred into the binary vector pGWB415 (Nakagawa et al., 2009) by Gateway LR reaction to create the final iFBW2 (FBW2 coding sequence including an intron) N-terminal 3HA fusion placed under the regulation of the 35S promoter.

The pUBQ:RFP-iFBW2 (also called pUBQ:FBW2) construct was obtained by Gateway LR recombination (Invitrogen) using pENTRY(221)-iFBW2 described earlier and the binary vector pUBN-RFP (Grefen et al., 2010) by Gateway LR reaction to create the final iFBW2 N-terminal RFP fusion placed under the regulation of the UBI10 promoter

For yeast two hybrid interaction assays, pGADT7-FBW2, pGBKT7-AGO1-NT-PAZ and pGBKT7-AGO1-L2-CT were generated. The *FBW2* coding sequence as well as AGO1 NT-PAZ and AGO1 L2-CT were amplified by PCR using primers listed in Table S3 and used for gateway recombination using pDONR-Zeo plasmid (Invitrogen). Then, pENTRY(Zeo)- FBW2, pENTRY(Zeo)-AGO1-NT-PAZ and pENTRY(Zeo)-AGO1-L2-CT were transferred into the pGADT7GW or pGBKT7GW by Gateway LR reaction to create pGADT7-FBW2, pGBKT7-AGO1-NT-PAZ and pGBKT7-AGO1-L2-CT. The pGADT7-ASK1 was described in (Pazhouhandeh et al., 2006).

To generate Gateway-compatible BIFC destination vectors, the rfA recombination cassette (Invitrogen) from the pGREEN-IIS binary destination vector pFK387 (Immink et al., 2012) was PCR-amplified and cloned as an HindIII/SmaI fragment into pSAT1-cEYFP- C1(B) and pSAT1-nEYFP-C1 BIFC vectors (Citovsky et al., 2006). The coding sequences of BPM5, SDE3, FBW2, AGO1 and AGO1-L2-CT were moved from the described ENTRY clones (Table S3) into the Gateway-compatible BIFC vectors by Gateway LR clonase reaction.

### Transient expression in *N. benthamiana* leaves and GFP fluorescence quantification

Agrobacterium cells (GV3101 Pmp90 or C58C1) harbouring the constructs of interest were grown overnight at 28°C in 10mL LB medium supplemented with antibiotics, resuspended in 10mM MgCl2 supplemented with 200mM acetosyringone at an OD of 0.3 per construct (unless otherwise specified), and incubated for 1 hour at room temperature before being pressure infiltrated into leaves of 4 week-old plants. Unless otherwise specified, all agro- infiltration assays were conducted in presence of P19. Plants were maintained in growth chambers under 16 hours light and 8 hours dark photoperiod with a constant temperature of 22°C. Sampling and observations were performed 72 hours after agro-infiltration.

GFP fluorescence emitted from the *N. benthamiana* agro-infiltrated leaves was quantified with a Ettan DIGE image (GE healthcare) with the parameters set for the SYPRO Ruby 1 dye (excitation filter 480/30 and emission filter 595/25) with 0.017 second exposure time.

### S-PTGS assay

GUS activity was quantified as described before (Gy et al., 2007) using crude extracts from plant leaves and monitoring the quantity of 4-methylumbelliferone products generated from the substrate 4-methylumbelliferyl-β-D-glucuronide (Duchefa) on a fluorometer (Thermo Scientific fluoroskan ascent).

### Confocal microscopy and Bimolecular Fluorescence Complementation (BIFC) assays

Confocal microscopy was performed on a LEICA TCS SP8 laser scanning microscope (Leica Microsystem) using the objective HCX APO CS 20X magnification with a numeric aperture of 0,7 without immersion. Usual excitation/detection-range parameters for CFP and Venus were 458 nm/465-510 nm and 514 nm/600-630 nm, respectively and emissions were collected using system hybrid (Hyd) detectors.

For the BiFC assays, isolation of Arabidopsis cell suspension protoplasts and PEG-mediated transformation was performed as described (Lalande et al., 2020). For each transformation, a combination of plasmid DNA (15μg) containing equal amounts of transfection control CPRF2-CFP (Bortolamiol et al., 2007) and each of the BIFC constructs was mixed with approximately 2 × 105 protoplasts. Protoplasts were cultured for 16 hours in the dark prior to confocal laser-scanning microscope observation. Images were taken under a Leica TCS SP8 microscope using CFP and YFP settings. The average value of YFP intensity per area of transfected protoplasts was quantified using Fiji software (ImageJ; http://imageJ.nih.gov/ij) and statistical significance of values between samples was calculated using the Student’s t-test method.

### Yeast two-hybrid assays

pGBKT7-AGO1-NT-PAZ (corresponding to AGO1 N-terminal part), pGBKT7-AGO1-L2- CT (corresponding to AGO1 C-terminal part), pGBKT7-ASK1 and pGADT7-FBW2 as well as pGBKT7, and pGADT7 empty vectors were used in yeast two-hybrid assays. The yeast strain PJ69-4 was transformed with the appropriate combinations of bait and prey vectors. Transformants were selected on synthetic (SD)/-Leu/-Trp (-LW) media (Clontech) and interactions assays were scored on selective medium deprived of Histidine (SD)/-Leu/-Trp/- His (-LWH) or Histidine and Adenine (SD)/-Leu/-Trp/-His/-Ade (-LWHA), allowing growth for 4 to 15 days at 28°C. Yeast transformation as well as yeast protein extraction were performed following the recommendations presented in the Yeast Protocol Handbook PT3024-1 (Clontech).

### Protein analysis and western blotting

Proteins were extracted in pre-heated (95°C) 2X Laemmli sample buffer, quantified using amido-black staining (Popov et al., 1975) and 10 to 20µg of total proteins were separated by SDS-PAGE, either on 7-12% Tris-glycine gels or gradient NuPAGE 4-12% Bis-Tris Protein Gels (Thermo Fischer) or gradient Criterion TGX gel (4-15%) (BioRad). List of antibodies and their working dilution used in this work are reported in (Table S4). For all western blots, immuno-luminescence was detected using the ECL Prime kit (GE Healthcare) or ECL Clarity (BioRad) and imaged using Fusion FX (Vilbert).

### Protein immuno-precipitation assays

For immunoprecipitation of HA-FBW2, 1g of frozen plant material (10 day-old seedlings) ground to a fine powder with a mortar and pestle, resuspended in 3 volumes of IP Extraction Buffer (25mM Tris HCl, pH 7.5, 150mM NaCl, 10% glycerol, 5mM MgCl2, 0.1% Tween20, 15mM EGTA, 10μM MG132, and 1× cOmplete™ Protease Inhibitor Cocktail [Roche]) and incubated for 30 min at 8 rpm in the cold room. Insoluble material was removed by centrifugation (twice 15 min, 16 000g, 4°C). Identical amounts of crude extracts were incubated with 25μl anti-HA magnetic beads (Pierce Anti-HA magnetic Beads) (pre-washed three times in IP Extraction buffer) for 3 hours at 8 rpm at room temperature. Immune complexes were washed three times in the IP extraction buffer. Elution of the immunoprecipitated proteins was performed by adding 30μl of glycine (0,2M pH 3) to the magnetic beads and transfer to a solution containing 10μl Tris HCl 1M pH 11. Before analysis on SDS-PAGE gels, 4X Laemmli loading buffer was added to a final concentration of 1X to the samples and then denatured for 5 minutes at 95°C.

For immunoprecipitation of endogenous AGO1, 500mg of frozen tissues (from 7 day-old seedlings) was ground to a fine powder with a mortar and pestle, resuspended in 3 volumes of crude extract buffer (50mM Tris, pH 7.5, 150mM NaCl, 10% glycerol, 5mM MgCl2, 0.1% IGEPALl, 5mM DTT, and 1x cOmplete™ Protease Inhibitor Cocktail [Roche]), and incubated for 20 minutes at 8 rpm in the cold room. Insoluble material was removed by centrifugation (twice 15 minutes, 16,000g, 4°C). Identical amounts of crude extracts were incubated with prebound @AGO1 (5μg) PureProteome Protein A magnetic beads (30μL; Millipore) for 2 hours at 7 rpm in the cold room. Immune complexes were washed four times in the crude extract buffer, and purified small RNA was eluted from the beads in Tri-Reagent (Sigma-Aldrich) following the manufacturer’s instructions.

### FBW2-immunoprecipitation, mass spectrometry analysis, data processing and availability

For each IP, 1g of seedlings was ground in liquid nitrogen for 10 minutes in 3 ml of ice-cold lysis buffer (50mM Tris, 50mM NaCl, 0.25% IGEPAL CA-630, 2mM MgCl2, 1mM DTT, 0.375% formaldehyde, protease inhibitors (cOmplete™–EDTA free, Roche). The crosslinked protein extract was quenched 2 minutes with glycine to a final concentration of 200mM. The cleared supernatants were divided in two affinity purifications, incubated with magnetic microbeads coupled to HA antibodies (Miltenyi, catalogue number 130-091-122), and complexes were eluted in 100 µl of pre-warmed elution buffer (Miltenyi). Co-IP experiments were performed in two independent biological replicates with two different transgenic lines (FBW2OE and FBW2OE/*ago1.27*. Each biological replicate was divided into two affinity-purification replicates. In parallel control IPs were carried out with HA antibodies in Col-0.

Eluted proteins were digested with sequencing-grade trypsin (Promega, Fitchburg, MA, USA). Each sample was further analyzed by nanoLC-MS/MS on a QExactive+ mass spectrometer coupled to an EASY-nanoLC-1000 (Thermo-Fisher Scientific, USA) as described previously (Chico et al., 2020). Data were searched against the TAIRv10 fasta protein sequences from *Arabidopsis thaliana* with a decoy strategy (27.282 forward protein sequences). Peptides and proteins were identified with Mascot algorithm (version 2.6.2, Matrix Science, London, UK) and data were further imported into Proline v2.0 software (http://proline.profiproteomics.fr/). Proteins were validated on Mascot pretty rank equal to 1, and 1% FDR on both peptide spectrum matches (PSM score) and protein sets (Protein Set score). The total number of MS/MS fragmentation spectra was used to quantify each protein from at least six independent biological and affinity replicates. After a DEseq2 normalization of the data matrix, the spectral count values were submitted to a negative-binomial test using an edgeR GLM regression through R (R v3.2.5). For each identified protein, an adjusted p-value (adjp) corrected by Benjamini–Hochberg was calculated, as well as a protein fold-change (FC). The results are presented in a Volcano plot using protein log2 fold changes and their corresponding adjusted (-log10adjp) to highlight upregulated and downregulated proteins. The mass spectrometry proteomics data have been deposited to the ProteomeXchange Consortium via the PRIDE [(Perez-Riverol et al., 2019)] partner repository with the dataset identifier PXD024840 and 10.6019/PXD024840.

### Microsomal fractionation

The crude cell extracts were prepared from 7 day-old seedlings that were ground with mortar and pestle in an ice-cold buffer containing 50mM HEPES pH 7.6, 30mM KCl, 5mM MgCl2, 5mM EGTA pH 8.0 and 250mM sucrose supplemented with freshly added 1mM DTT and Protease Inhibitor cocktail (Roche). After centrifugation at 1,000 x g for 5 minutes at 4°C the resulting supernatant represents the total extract (Total). The microsomal (Micro) and cytoplasmic (Cyto) fractions were collected by centrifugation of soluble cell extract in a TLA- 110 rotor (Beckman Coulter ultracentrifuge) at 100,000 x g for 30 minutes at 4°C. For protein analysis, fractions were precipitated with methanol/chloroform and protein pellets were dissolved in 1x Laemmli buffer.

### Size exclusion chromatography

About 800mg seedlings of the indicated genotypes were extracted in 2.5 Volumes of 50mM Tris-HCl pH7.5, 150mM NaCl, 5mM MgCl2, 10% glycerol, 0.1% IGEPAL, 5mM DTT, 10µM MG132, 1X cOmplete™ Protease Inhibitor Cocktail (Roche) and left to mix on carousel for 30 minutes. Extracts were centrifuged 10 minutes at 4400 rpm in Falcon tubes, filtered through Miracloth, and filtered through a 0,2µm Minisart RC 4 syringe filter. Crude extracts were calibrated to 1.9µg/µl using the amidoblack method, and 500µl were injected in 4 separate loops. Separation was performed sequentially on a Superdex 200 10/300 increase column on an AKTA Pure system with the following settings: 500µl/min, fraction volume of 250µl collected from 7.25ml to 13ml. For proteins, half of the fractions was precipitated in 2 volumes of absolute ethanol at 4°C for 48 hours. Samples were centrifuged at maximum speed for 30 minutes at 4°C, pellets were resuspended in 2x Laemmli buffer and treated at 95°C for 5 minutes. Denatured samples were separated in 7.5% acrylamide SDS PAGE gels and treated as indicated in the immunoblot section. Input samples were collected from the crude extract and analysed separately following the same method. For RNA, half the fraction (every two fraction) was mixed with 300µl tri-Reagent and extracted as indicated. Precipitated RNA was resuspended in a final concentration of 60% formamide, 5mM EDTA, 0.05% bromophenol blue-0.05% xylene cyanol, heated a 95°C for 5 minutes, and separated by electrophoresis on a 15% polyacrylamide (19:1 acrylamide:bisacrylamide), 8M Urea, 0.5× TBE gel at 15 watts. Electroblotting, crosslinking and hybridization was performed as standard. Input samples were collected from the original plant material and analysed separately following the same method.

### Radiolabelling of AGO1 co-purified RNA

AGO1 immunoprecipitation were performed as indicated, from 400mg of 2 week-old seedlings of the indicated genotypes. Purified RNA was eluted from the beads in tri-Reagent, as indicated, and RNA was precipitated overnight at -20°C in 50% isopropanol and 40µg of glycogen. Stabilization of AGO1 in the *hyl1-2/fbw2-4* crude extract was verified by immunoblot before proceeding with labelling. Precipitated RNA was resuspended in 10µl of ultrapure water and 5µl was [γ-32P]ATP labelled by T4 polynucleotide kinase (Thermo Fisher Scientific) for 35 min at 37°C in buffer B. Labelled RNA was Tri-Reagent extracted as indicated, the aqueous phase filtered through a G25 MicroSpin column (GE healthcare) and the flowthrough precipitated in 75% isopropanol with 40µg of glycogen overnight at −20°C. A control reaction with 90ng of siR255 (21-nt) RNA oligo was treated in an identical fashion. Labelled RNA was resuspended in a final concentration of 60% formamide-5 mM EDTA- 0.05% bromophenol blue-0.05% xylene cyanol, heated a 95°C for 5 min, and separated by electrophoresis on a 15% polyacrylamide (19:1 acrylamide:bisacrylamide), 8M urea, 0.5×TBE gel at 15 watts for 230 minutes. The gel was wrapped in plastic and the signal was detected using FUJI medical x-ray films.

### GW motif prediction

GW motif prediction tool from http://www.combio.pl/agos/help/ (Karlowski et al., 2010)(Zielezinski and Karlowski, 2011) identified a putative AGO hook motif in the FBW2 protein sequence from residue 287 to 299.

### RT-qPCR and small RNA analyses by northern blotting

For quantitative RT-PCR (qPCR), 1μg of total RNA extracted in Tri-Reagent according to the manufacturer’s instruction was treated with DNaseI (Fisher Scientific) and reverse transcribed with High-Capacity cDNA Reverse Transcription Kit (Applied Biosystem). PCR was performed using gene specific primers (see Table S3) in a total volume of 10μl SYBR Green Master mix (Roche) on a LightCycler LC480 apparatus (Roche) according to the manufacturer’s instructions. The mean value of three replicates was normalized using the EXP (AT4G26410), and TIP41 (AT4G34270) genes as internal controls. All primers used in qRT-PCR are listed in Table S3.

For small RNA analysis, RNA was extracted in Tri-Reagent according to the manufacturer’s protocol, and aqueous phase was left in 1 volume of isopropanol over-night at -20°C, precipitated 30 minutes at 16000g (4°C). Pellets were rinsed in 1ml 70% ethanol and centrifuged an additional 10 minutes. Dry RNA pellets were resuspended in 60% deionised formamide. RNA gel blot, transfer, crosslinking and hybridization were performed as described in (Derrien et al., 2018), with 10μg of RNA loaded in each lane. DNA oligonucleotides complementary to miR403, miR168, miR159 and U6 RNA (see Table S3) were 5’ end-labelled with [γ-32P]ATP using T4 polynucleotide kinase (PNK) (Promega).

### Libraries preparation and high-throughput sequencing

Total RNA samples were extracted from 1-week-old Col-0, *hyl1-2*, *hyl1-2 fbw2-4*, *hen1-6*, *hen1-6 fbw2-4* and *fbw2-4* seedlings grown on MS-agar plates using Tri-Reagent according to the manufacturer’s instruction. For AGO1-loaded small RNA samples, IPs were performed as described above from 500mg of 1-week-old Arabidopsis seedlings grown on MS-agar plates. AGO1-loaded small RNA were then extracted by adding Tri-Reagent directly on the magnetic beads and extraction of RNA was then performed according to the manufacturer’s instructions.

Small RNA libraries were constructed using Real-Seq-AC kit (RealSeq®-AC, USA) according to manufacturer’s instructions with 500ng as starting material. Libraries were sequenced using Illumina Next-Seq 500 technology at University of Delaware (Delaware, USA). Parallel analysis of RNA end (PARE) libraries were constructed following the previously published protocol (Zhai et al., 2014) using 20ug of total RNA as starting material, and sequenced using an Illumina HiSeq 2500 instrumenty at the University of Vermont (Vermont, USA).

### Sequencing data analysis and availability

For this study, a total of 36 small RNA libraries (with at least 10 M mapped reads were ultimately used) and 12 PARE libraries were constructed. For small RNA libraries, we trimmed adapters and low-quality reads using Cutadapt v2.9 software (Martin, 2011), retaining only reads between 18- and 34-nt in length. Reads were then mapped to the Arabidopsis genome version 10 (available at www.arabidopsis.org/download/) and its corresponding TAIR10 BLAST sets for all the features, using Bowtie2 (Langmead and Salzberg, 2012). Differential accumulation was done using DESeq2 (Love et al., 2014) and all plots were generated using ggplot2 (Wickham, 2016) packages in R environment. PARE libraries were trimmed and quality checked using the same tools as small RNA libraires, and analysed using CleaveLand v4.5 (Addo-Quaye et al., 2009). To achieve a higher reliability, we only considered small RNA/target signatures in categories 0-1-2, that are present in all the biological replicates, and produced by small RNA identified in AGO1 IP libraries between 19 and 24 nt in length. Gene Ontology (GO) analysis was performed using PlantRegMap tool (Tian et al., 2020) and the GO term enrichment tool with all the identified genes as background and p-value Bonferroni correction.

The data discussed in this publication have been deposited in NCBI’s Gene Expression Omnibus (Edgar et al., 2002) and are accessible through GEO Series accession number GSE169324 for small RNA seq (https://www.ncbi.nlm.nih.gov/geo/query/acc.cgi?acc=GSE169324) and GSE169434 for PARE seq (https://www.ncbi.nlm.nih.gov/geo/query/acc.cgi?acc=GSE169434).

## Acknowledgments

We thank Scott Poethig for sharing material and a fruitful discussion. Nicolas Baumberger for help with gel filtration assays. P.G. acknowledges support from the European Research Council under the European Union’s Seventh Framework Programme (FP7/2007-2013) / ERC advanced grant to PG agreement n° [338904] and ANR-10-LABX-0036_NETRNA and ANR-17-EURE-0023 programs managed by the French National Research Agency. The mass spectrometry instrumentation was funded by the University of Strasbourg, IdEx “Equipement mi-lourd” 2015. Work in the Meyers lab was supported by US National Science Foundation IOS award 1842685 and resources from the Donald Danforth Plant Science Center and the University of Missouri.

## Author contributions

TH, MC, and PG designed research; TH, MC, PB, EL, IPS, MS, BD, MD, PH, LK, DB, NB, HV performed research; TH, MC, PB, HV, BM and PG analysed the data; PG wrote the paper, with help from TH.

