## Supplementary material for "The Arabidopsis F-box protein FBW2 degrades AGO1 to avoid spurious loading of illegitimate small RNA": Supp Files

**Supporting Information**

**Supp Fig Legends**

**FIGURE S1: Degradation by FBW2 of the AGO1 protein domains.**

(A) Schematic representation of Arabidopsis AGO1 color-coded protein domains: DUF1785: Domain of Unknown Function 1785. PAZ: Piwi-Argonaute-Zwille. L2: Linker 2. MID: Middle domain.

(B-C) Western blots of protein extracts from 4 week-old *N. benthamiana* agroinfiltrated leaves. Agrobacteria harbouring binary vectors for the expression of 35S:3HA-FBW2, 35S:CFP-AGO1 or AGO1 protein domains C-terminally fused to the GFP, were infiltrated at 0.3 OD and tissues were sampled 3 days later. Expression of 35S:GUS serves as control. Coomassie blue (CB) staining was used as a loading control. The "@" symbol indicates hybridization with the corresponding antibodies.

**FIGURE S2: Overexpression of 3HA-FBW2 *in planta* and phenotypic characterization of the mutants.**

(**A**) Western blot of protein extracts from 4 week-old *N. benthamiana* agro-infiltrated leaves. Agrobacteria harbouring binary vectors for the expression of 35S:CFP-AGO1 combined with either 35S:GUS (control), 35S:P0-6myc, 35S:FBW2 and 35S:3HA-FBW2 constructs were infiltrated at an OD of 0,3 and tissues were sampled 3 days later. Coomassie blue (CB) staining was used as loading control. The "@" symbol indicates hybridization with the corresponding antibody.

(**B**) Three representative independent homozygous, simple insertion, FBW2OE (35S:3HA-FBW2) transgenic lines expressing variable amount of 3HA-FBW2 protein. Protein extracts from 7 day-old seedlings grown on MS medium were analysed by Western blot. Coomassie blue (CB) staining was used as loading control. The "@" symbol indicates hybridization with the corresponding antibodies.

(**C**) Average area of each individual leaf, from old to young (L1-L14), of *fbw2-1*, *fbw2-4* and the FBW2OE line 10. Measurements were performed at 22 days after stratification of plants grown in soil. Ten plants were measured for each line.

(**D**) Number of lateral roots of young seedlings of *fbw2-4* and FBW2OE line 10. Lateral roots were counted at 8 days after stratification after growth *in vitro* on vertical petri dishes. Lateral roots were counted under a binocular (lateral root stage >VIII) for at least 10 plants per line. *** = $P<0.001$, t-test.

**FIGURE S3: Inducible expression of 3HA-FBW2 in wild type and PTGS-deficient Arabidopsis mutant lines.**

(**A-B**) Kinetic analysis of transgenic *Arabidopsis* Col-0 XVE:3HA-FBW2 (A) and Col-0 XVE-P0-6myc (B) lines. Western blots of protein extracts from 5 to 12 day-old seedlings grown on MS medium supplemented with DMSO (-) or β-Es (10$\mu$M) (+). Coomassie blue (CB) staining was used as loading control and the "@" symbol indicates hybridization with the corresponding antibodies.

(**C**) Western blot of protein extracts from 7 day-old seedlings XVE-P0-myc and XVE:3HA-FBW2 crossed with the specified *ago1* mutants and grown on MS medium supplemented with DMSO (-) or β-Es (10$\mu$M) (+). Coomassie blue (CB) staining was used as loading control and the "@" symbol indicates hybridization with the corresponding antibodies.
(**D**) FBW2 expression in the *ago1-57* mutant background. Upper panel: Kinetic analysis of transgenic Arabidopsis XVE:3HA-FBW2/*ago1-57* line. Western blots of protein extracts from 5 to 12 day-old seedlings grown on MS medium supplemented with DMSO (-) or β-Es (10$\mu$M) (+). Coomassie blue (CB) staining was used as loading control and the "@" symbol indicates hybridization with the corresponding antibodies. Bottom panel: RT-qPCR analysis of *FBW2* expression level, relative to the 5 day-old Col-0, in seedlings of the indicated genotypes grown on MS medium supplemented with β-Es (10$\mu$M).
(**E**) Kinetic analysis of transgenic *Arabidopsis* XVE:3HA-FBW2/*sgs2-1* line. Western blots of protein extracts from 5 to 11 day-old seedlings grown on MS medium supplemented with DMSO (-) or β-Es (10$\mu$M) (+). The arrow represents the AGO2 protein band while * represents the remaining AGO1 signal from the previous hybridization. Coomassie blue (CB) staining was used as loading control and the "@" symbol indicates hybridization with the corresponding antibodies.

**FIGURE S4: Assessment of protein expression for the yeast two-hybrid interaction assays.** Western blot of protein extracts from yeast parental lines. Coomassie blue (CB) staining was used as a loading control and the "@" symbol indicates hybridization with the corresponding antibodies. Expected sizes for the proteins are 77 kDa for AGO1 NT-PAZ, 83 kDa for AGO1 L2-CT, 40 kDa for ASK1 and 56 kDa for FBW2.

**FIGURE S5: Effect of the inverted repeat *gffg* and P19 overexpression on AGO1 degradation by FBW2.**
(**A**) Western blot of protein extracts from 4 week-old *N. benthamiana* agroinfiltrated leaves with Agrobacteria harbouring the following binary vectors: 35S:CFP-AGO1, 35S:GUS (control), 35S:3HA-FWB2, 35S:gffg and 35S:P19 in combinations as specified. All constructs were infiltrated at 0,3 OD and tissues for protein analysis were sampled 3 days later. Coomassie blue (CB) staining was used as loading control. For this panel and also for panel B, AGO1 signal was quantified by ImageJ, normalized to the corresponding CB. Numbers are indicated below the panel as relative to the control set at 1.0. The "*" symbol indicates an aspecific cross-reacting band. The "@" symbol indicates hybridization with the corresponding antibodies.
(**B**) Western blot of protein extracts from 4 week-old *N. benthamiana* agroinfiltrated leaves with Agrobacteria harbouring the following binary vectors: 35S:RFP-AGO1, 35S:GUS (control), 35S:RFP-FWB2, 35S:gffg and 35S:P19 in combinations as specified. All constructs were infiltrated at 0,2 OD except for the experiment with P19 for which the infiltration solutions were adjusted to compensate the difference of expression (0.05 for AGO1, 0.1 for GUS and 0.1 for FBW2). Coomassie blue (CB) staining was used as loading control. The "*" symbol indicates an aspecific cross-reacting band. The "@" symbol indicates hybridization with the corresponding antibodies.
(**C**) Kinetic analysis of transgenic Arabidopsis XVE:3HA-FBW2/ *fbw2*-1/35S:3HA-P19 line. Western blots of protein extracts from 5 to 11 day-old seedlings grown on MS medium

supplemented with DMSO (-) or β-Es (10$\mu$M) (+). Coomassie blue (CB) staining and ACTIN protein levels were used as loading controls. The "@" symbol indicates hybridization with the corresponding antibodies. The "*" symbol represents the remaining AGO1 signal from the previous hybridization.

**FIGURE S6: AGO1 protein level and plant phenotype of *hen1-6 fbw2-4* and *hyl1-2 fbw2-4* double mutants.**

(**A**) Western blot of protein extracts from 5 day-old seedlings of the indicated genotypes. Hybridization with the CDC2 antibody and Coomassie blue (CB) staining were used as loading controls.

(**B**) Pictures of three representative plants per genotype just before bolting stage showing worsening of the phenotype in the double mutants.

**FIGURE S7. Loss of *FBW2* restores high molecular weight AGO1 complexes in *hyl1-2*.**

(**A**) Western blot of protein extracts of 13 day-old seedlings of Col-0, *fbw2-4*, *hyl1-2*, and *hy1-2 fbw2-4*. Two biological replicates were made to illustrate the tendency of AGO1 homeostasis.

(**B**) Gel filtration analysis of AGO1-based RISC complexes from the same protein extracts shown in (A). Western blots of protein extracts from eluates of the gel filtration column (superdex 200 10/300 increase column on an AKTA Pure system) spanning from 6.75 to 13.75 ml elution volume. Molecular weights (kDa) of known protein sizes are indicated on top of the blot. "@" indicates hybridization with the corresponding antibodies. Coomassie blue staining was used as loading control.

**FIGURE S8. AGO1 protein level in different mutants used for RNA deep-sequencing experiments.**

Western blot of protein extracts from 8 day-old seedlings of the indicated genotypes. R#1, R#2, R#3 correspond to the three biological replicates. Hybridization with the @AGO1 antibody. Coomassie blue (CB) staining were used as loading controls

**FIGURE S9. *FBW2* mutation has no effect in small RNA size distribution.**

Size distribution of small RNA mapping to the Arabidopsis TAIR10 genome; the abundance of reads per million (RPM) of each size class was calculated for each source of data. The x axis indicates the small RNA size (from 18 to 34 nt) and the y axis indicates its abundance. Small RNA distribution for total RNA samples(A) and small RNA size distribution for AGO1 IP samples (B).

**FIGURE S10. *FBW2* mutation has a minimal effect in the miRNA population.**

Relative abundance of miRNA with significant differential expression in single and double mutants compared to wild-type (Col-0) and single mutants; the relative abundance is expressed as a heat map (see Key at the bottom), with the samples being compared indicated below each heat map (*Q value # 0.05, **Q value # 0.01 and **Q value #0.001).

**FIGURE S11. Pol4- and TE-derived sRNA reads are enriched in *hyl1 fbw2* double mutant, while no significant differences are observed in *fbw2* single mutants.**

Boxplot representing the abundance of reads (in reads per million) mapping to eight different features of the Arabidopsis genome TAIR 10 for total RNA samples. These include the following from left to right: cDNA; mature miRNA; siRNA precursors dependent on Pol4; ribosomal RNA (rRNA); small nuclear and small nucleolar RNA (snRNA and snoRNA); TAS precursors; Transposable elements (TEs); tRNA-derived sRNA (tRNA); RPM, reads per million. Three biological replicates are included for each sample.

**TABLE S1: Proteins identified in control and co-IP samples**

**TABLE S2: Deep-sequencing analyses of total and AGO1-IP small RNA and PARE.**

**TABLE S3: Table of genotyping, RT-qPCR and cloning primers and Northern probes.**

**TABLE S4 List of antibodies and chemicals used in this work.**

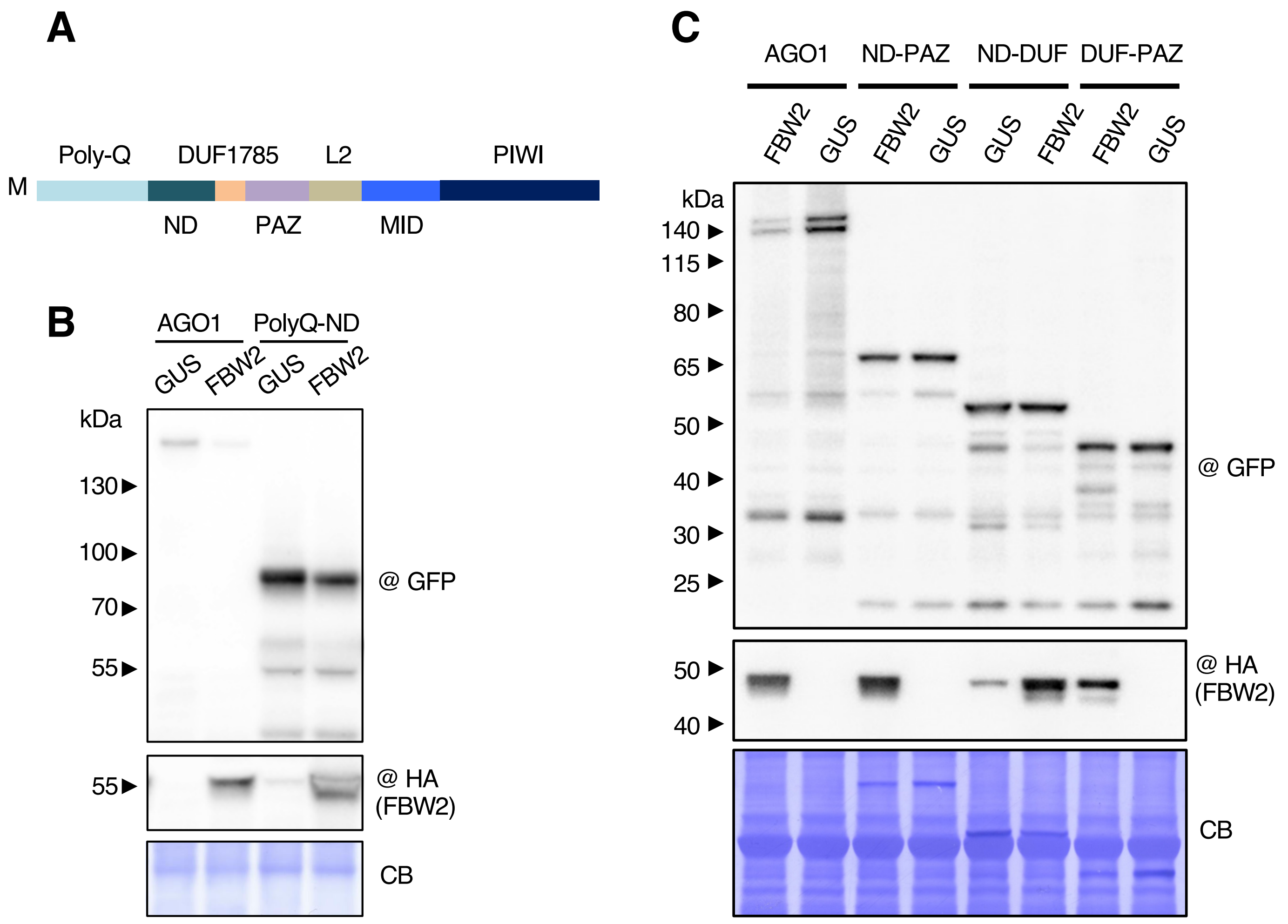
A
Poly-Q
DUF1785
L2
PIWI
M
ND
PAZ
MID
B
AGO1
PolyQ-ND
GUS
FBW2
GUS
FBW2
kDa
130
100
70
55
@ GFP
55
@ HA
(FBW2)
CB
C
AGO1
ND-PAZ
ND-DUF
DUF-PAZ
FBW2
GUS
FBW2
GUS
GUS
FBW2
FBW2
GUS
kDa
140
115
80
65
50
40
30
25
@ GFP
50
40
@ HA
(FBW2)
CB

**Figure S1**

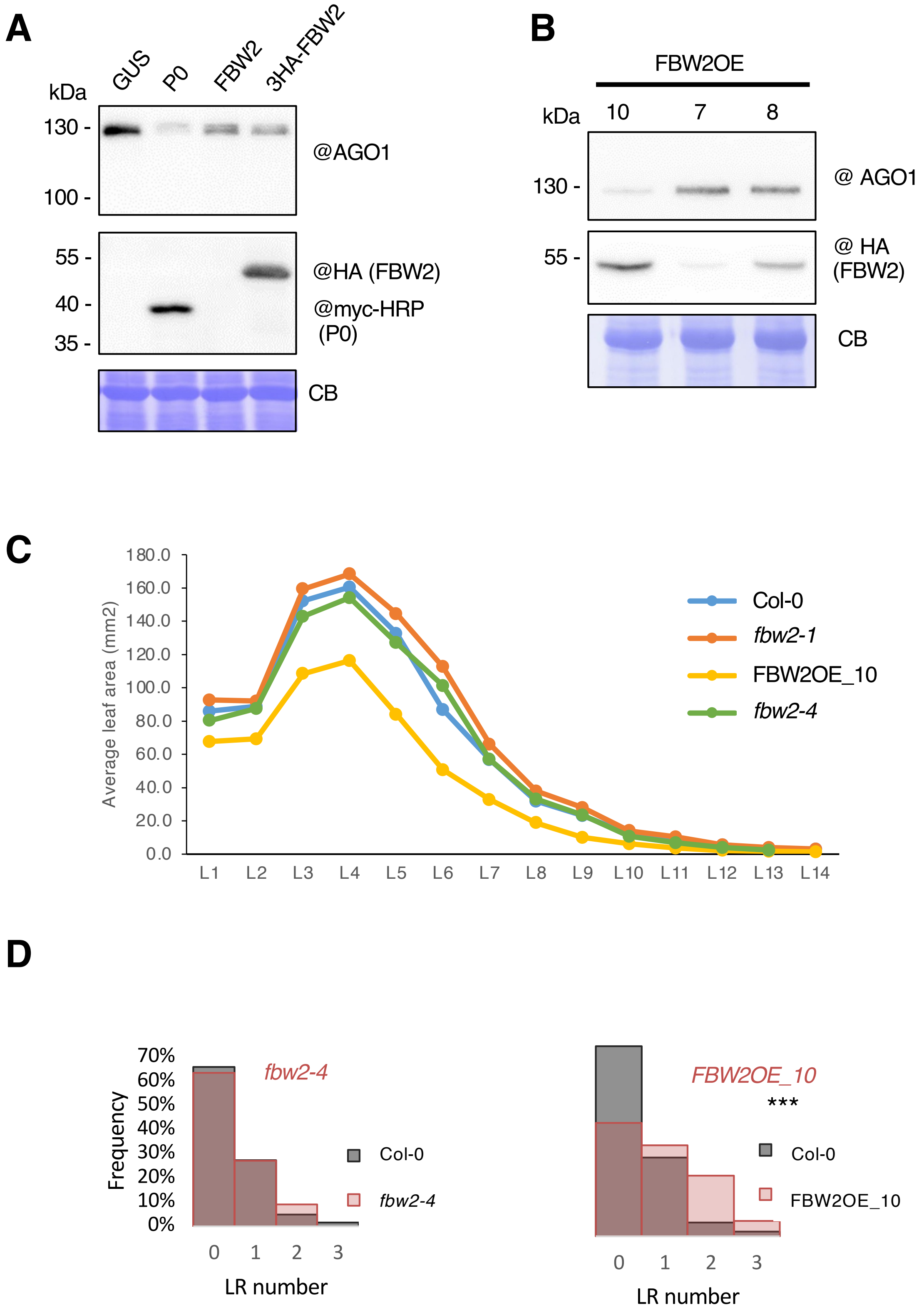
A
kDa
GUS
P0
FBW2
3HA-FBW2
130 -
100 -
@AGO1
55 -
40 -
35 -
@HA (FBW2)
@myc-HRP
(P0)
CB
B
FBW2OE
kDa
10
7
8
130 -
@ AGO1
55 -
@ HA
(FBW2)
CB
C
Average leaf area (mm2)
180.0
160.0
140.0
120.0
100.0
80.0
60.0
40.0
20.0
0.0
L1
L2
L3
L4
L5
L6
L7
L8
L9
L10
L11
L12
L13
L14
Col-0
fbw2-1
FBW2OE_10
fbw2-4
D
Frequency
70%
60%
50%
40%
30%
20%
10%
0%
fbw2-4
Col-0
fbw2-4
0
1
2
3
LR number
FBW2OE_10
***
Col-0
FBW2OE_10
0
1
2
3
LR number

**Figure S2**

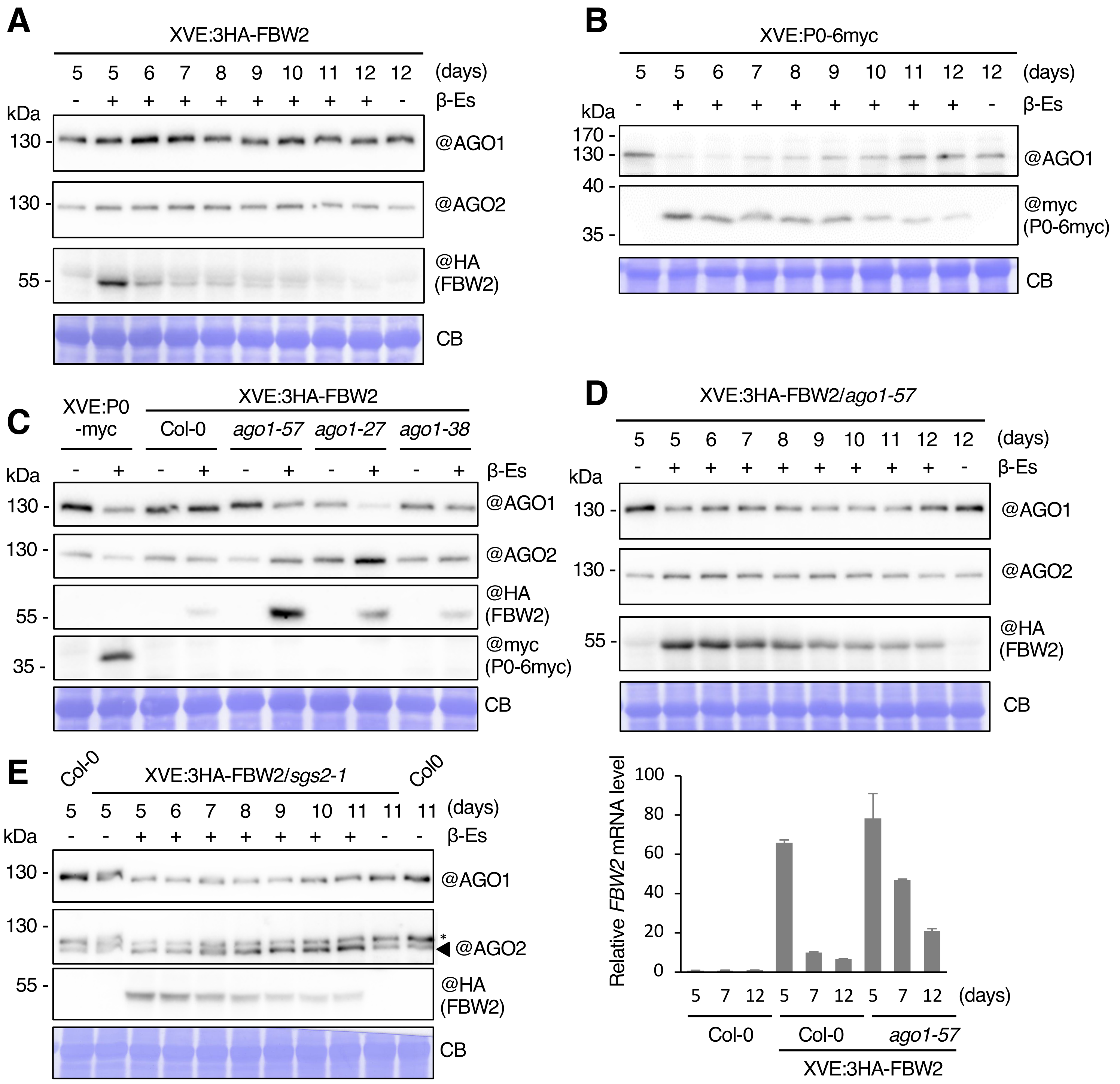

A
XVE:3HA-FBW2
5 5 6 7 8 9 10 11 12 12 (days)
- + + + + + + + + - β-Es
kDa
130 -
@AGO1
130 -
@AGO2
55 -
@HA (FBW2)
CB
B
XVE:P0-6myc
5 5 6 7 8 9 10 11 12 12 (days)
- + + + + + + + + - β-Es
kDa
170 -
130 -
@AGO1
40 -
35 -
@myc (P0-6myc)
CB
C
XVE:P0 -myc
XVE:3HA-FBW2
Col-0
ago1-57
ago1-27
ago1-38
- + - + - + - + - + β-Es
kDa
130 -
@AGO1
130 -
@AGO2
55 -
@HA (FBW2)
35 -
@myc (P0-6myc)
CB
D
XVE:3HA-FBW2/ago1-57
5 5 6 7 8 9 10 11 12 12 (days)
- + + + + + + + + - β-Es
kDa
130 -
@AGO1
130 -
@AGO2
55 -
@HA (FBW2)
CB
E
Col-0
XVE:3HA-FBW2/sgs2-1
Col0
5 5 5 6 7 8 9 10 11 11 11 (days)
- - + + + + + + + - - β-Es
kDa
130 -
@AGO1
130 -
*
@AGO2
55 -
@HA (FBW2)
CB
Relative FBW2 mRNA level
100
80
60
40
20
0
5 7 12 5 7 12 5 7 12 (days)
Col-0
Col-0
ago1-57
XVE:3HA-FBW2


### Figure S3

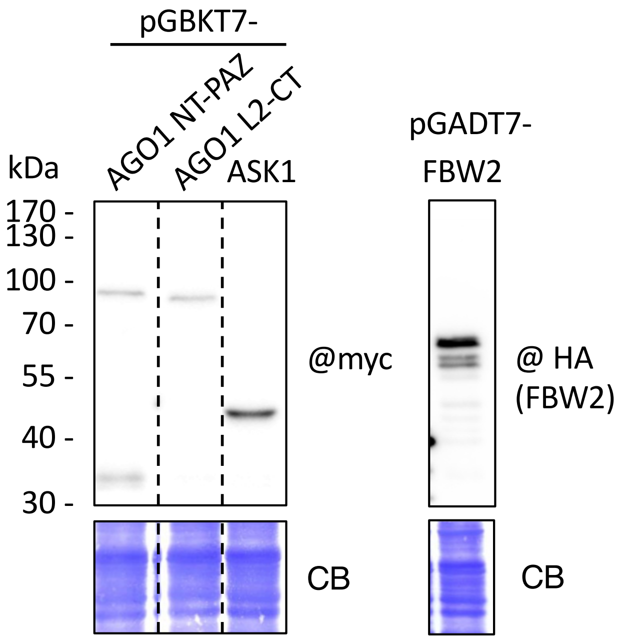
pGBKT7-
AGO1 NT-PAZ
AGO1 L2-CT
ASK1
pGADT7-
FBW2
kDa
170 -
130 -
100 -
70 -
55 -
40 -
30 -
@myc
@ HA
(FBW2)
CB
CB

**Figure S4**

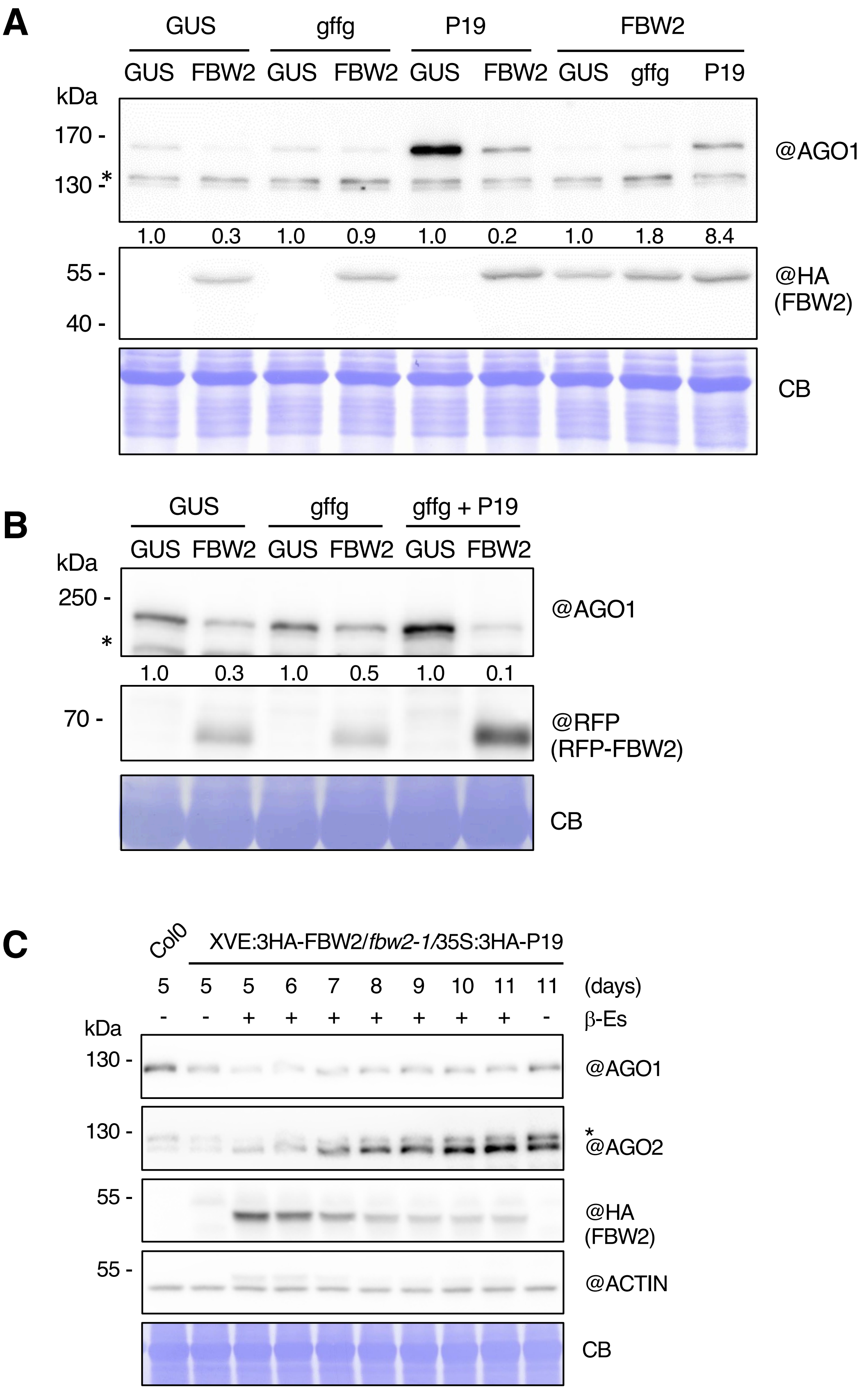
A
GUS
gffg
P19
FBW2
GUS FBW2 GUS FBW2 GUS FBW2 GUS gffg P19
kDa
170 -
130 -
*
@AGO1
1.0 0.3 1.0 0.9 1.0 0.2 1.0 1.8 8.4
55 -
40 -
@HA
(FBW2)
CB
B
GUS
gffg
gffg + P19
GUS FBW2 GUS FBW2 GUS FBW2
kDa
250 -
*
@AGO1
1.0 0.3 1.0 0.5 1.0 0.1
70 -
@RFP
(RFP-FBW2)
CB
C
Col0
XVE:3HA-FBW2/fbw2-1/35S:3HA-P19
5 5 5 6 7 8 9 10 11 11 (days)
- - + + + + + + + - β-Es
kDa
130 -
@AGO1
130 -
*
@AGO2
55 -
@HA
(FBW2)
55 -
@ACTIN
CB

**Figure S5**

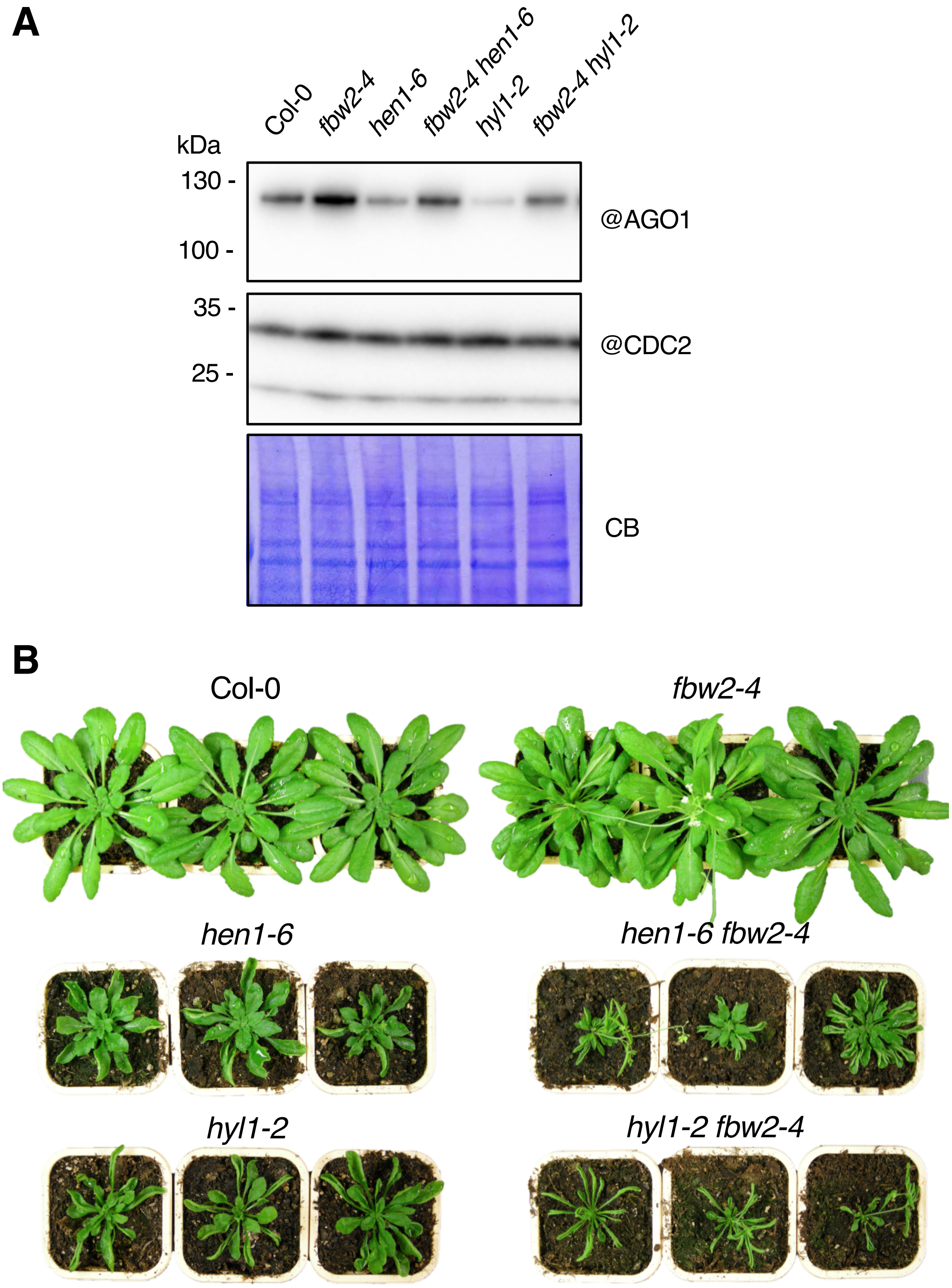
A
Col-0
fbw2-4
hen1-6
fbw2-4 hen1-6
hyl1-2
fbw2-4 hyl1-2
kDa
130 -
100 -
@AGO1
35 -
25 -
@CDC2
CB
B
Col-0
fbw2-4
hen1-6
hen1-6 fbw2-4
hyl1-2
hyl1-2 fbw2-4

**Figure S6**

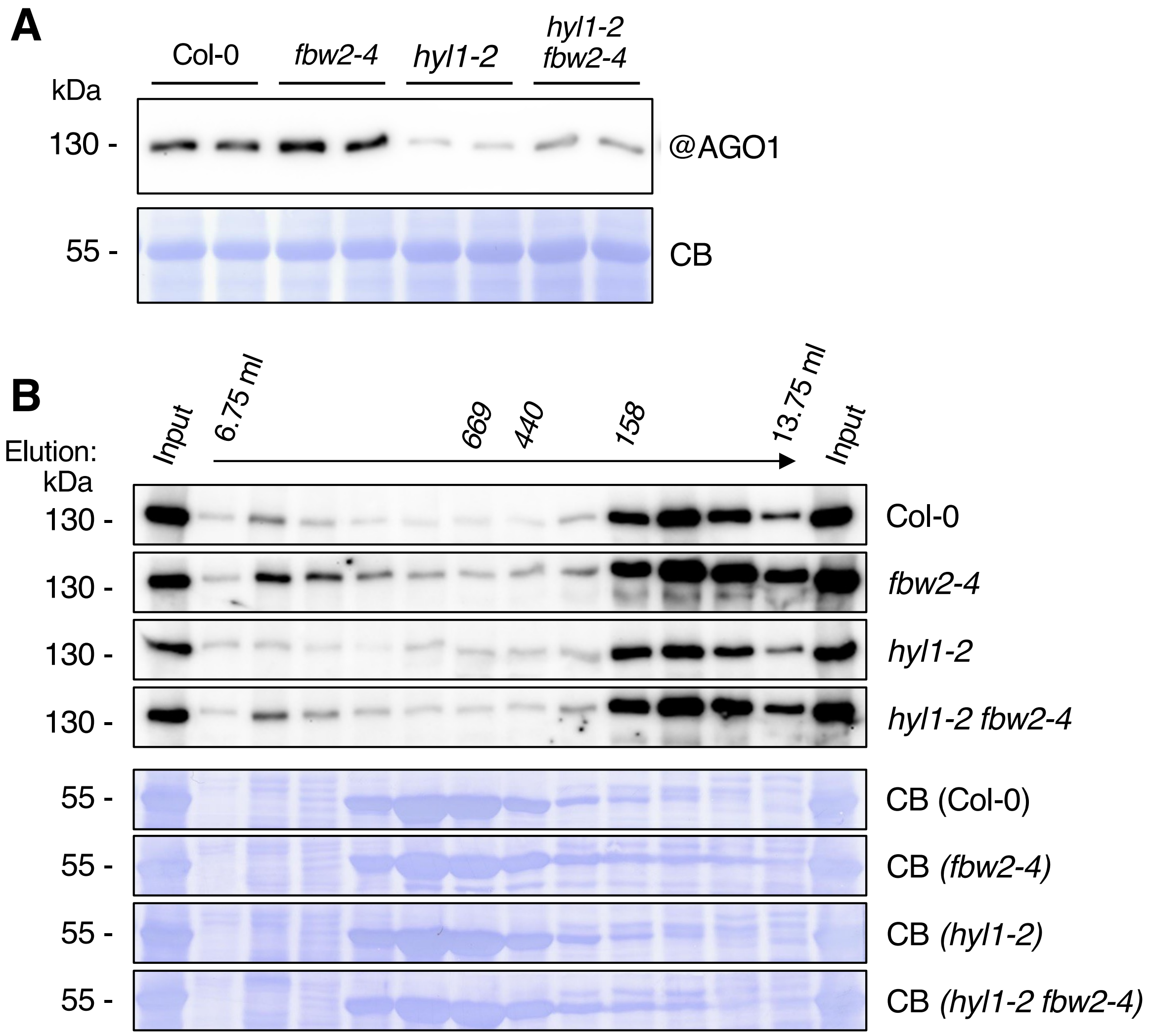
A
Col-0
fbw2-4
hyl1-2
hyl1-2 fbw2-4
kDa
130 -
@AGO1
55 -
CB
B
Elution:
Input
6.75 ml
669
440
158
13.75 ml
Input
kDa
130 -
Col-0
fbw2-4
hyl1-2
hyl1-2 fbw2-4
55 -
CB (Col-0)
CB (fbw2-4)
CB (hyl1-2)
CB (hyl1-2 fbw2-4)

**Figure S7**

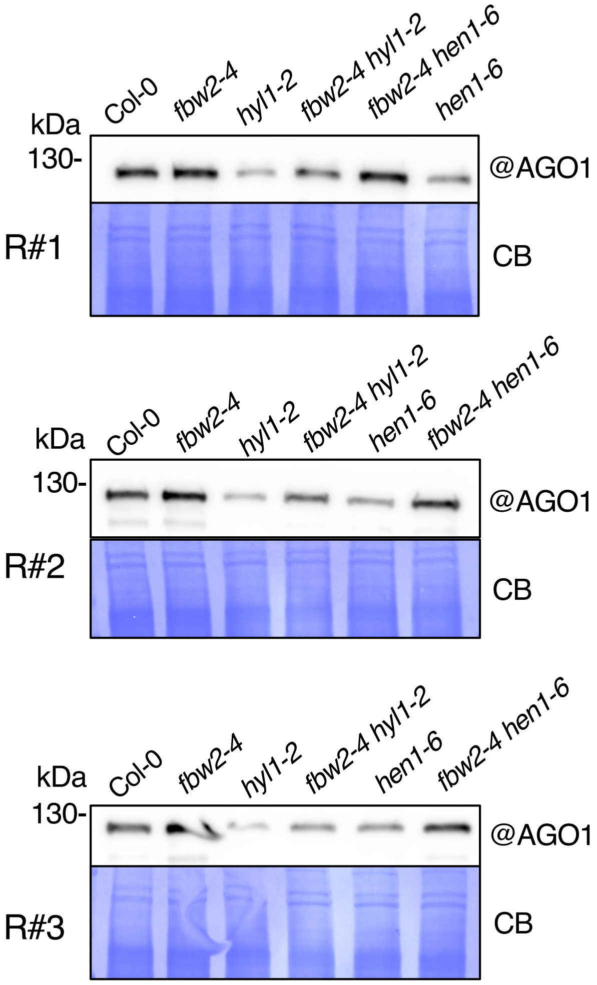
Col-0
fbw2-4
hyl1-2
fbw2-4 hyl1-2
fbw2-4 hen1-6
hen1-6
kDa
130-
@AGO1
R#1
CB
Col-0
fbw2-4
hyl1-2
fbw2-4 hyl1-2
hen1-6
fbw2-4 hen1-6
kDa
130-
@AGO1
R#2
CB
Col-0
fbw2-4
hyl1-2
fbw2-4 hyl1-2
hen1-6
fbw2-4 hen1-6
kDa
130-
@AGO1
R#3
CB

**Figure S8**

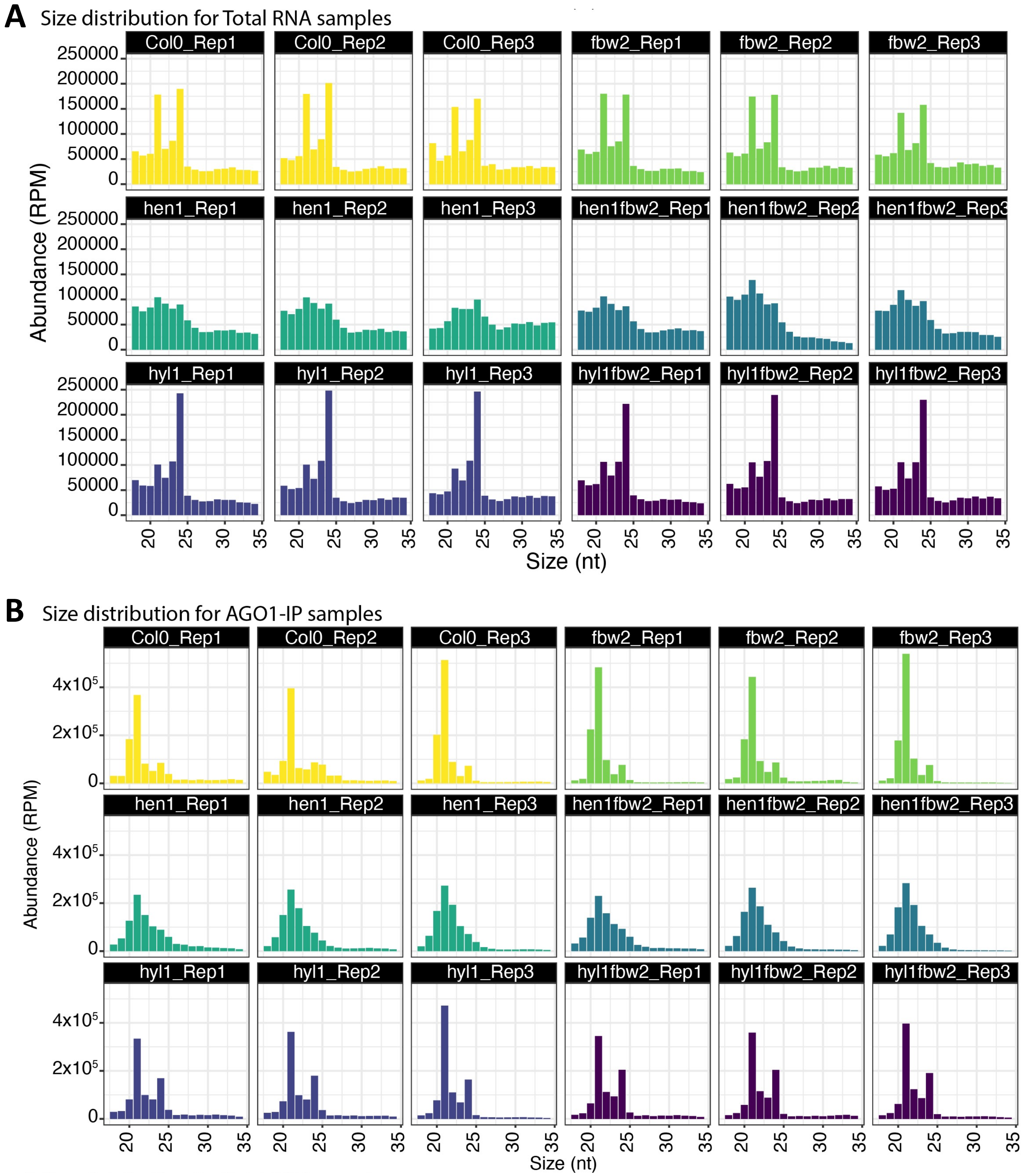
A
Size distribution for Total RNA samples
Col0_Rep1
Col0_Rep2
Col0_Rep3
fbw2_Rep1
fbw2_Rep2
fbw2_Rep3
hen1_Rep1
hen1_Rep2
hen1_Rep3
hen1fbw2_Rep1
hen1fbw2_Rep2
hen1fbw2_Rep3
hyl1_Rep1
hyl1_Rep2
hyl1_Rep3
hyl1fbw2_Rep1
hyl1fbw2_Rep2
hyl1fbw2_Rep3
250000
200000
150000
100000
50000
0
Abundance (RPM)
20 25 30 35
Size (nt)
B
Size distribution for AGO1-IP samples
Col0_Rep1
Col0_Rep2
Col0_Rep3
fbw2_Rep1
fbw2_Rep2
fbw2_Rep3
hen1_Rep1
hen1_Rep2
hen1_Rep3
hen1fbw2_Rep1
hen1fbw2_Rep2
hen1fbw2_Rep3
hyl1_Rep1
hyl1_Rep2
hyl1_Rep3
hyl1fbw2_Rep1
hyl1fbw2_Rep2
hyl1fbw2_Rep3
4x10⁵
2x10⁵
0
Abundance (RPM)
20 25 30 35
Size (nt)

**Figure S9**

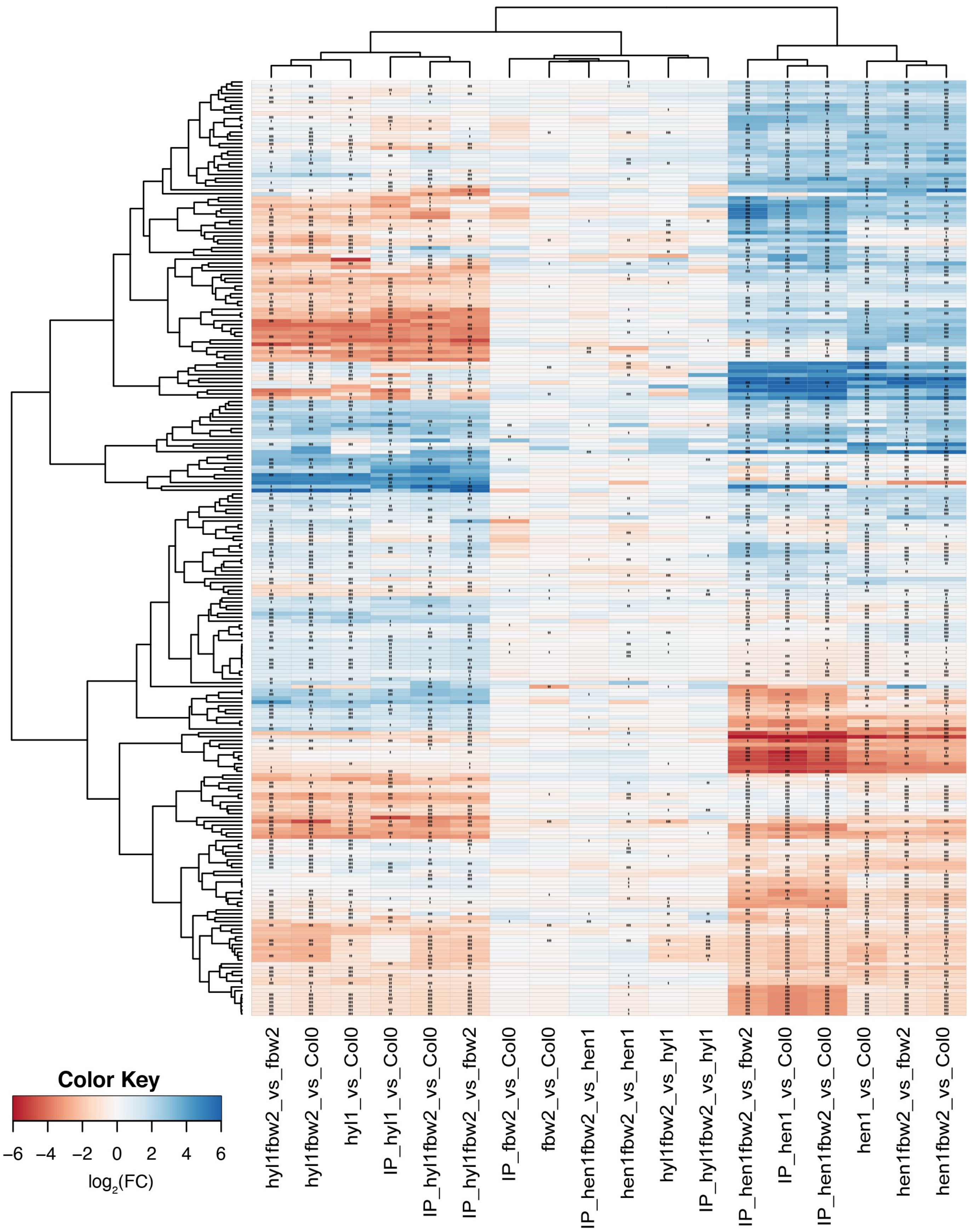
Color Key
−6 −4 −2 0 2 4 6
log2(FC)
hyl1fbw2_vs_fbw2
hyl1fbw2_vs_Col0
hyl1_vs_Col0
IP_hyl1_vs_Col0
IP_hyl1fbw2_vs_Col0
IP_hyl1fbw2_vs_fbw2
IP_fbw2_vs_Col0
fbw2_vs_Col0
IP_hen1fbw2_vs_hen1
hen1fbw2_vs_hen1
hyl1fbw2_vs_hyl1
IP_hyl1fbw2_vs_hyl1
IP_hen1fbw2_vs_fbw2
IP_hen1_vs_Col0
IP_hen1fbw2_vs_Col0
hen1_vs_Col0
hen1fbw2_vs_fbw2
hen1fbw2_vs_Col0

**Figure S10**

#### sRNA reads mapping Arabidopsis selected features for Total RNA

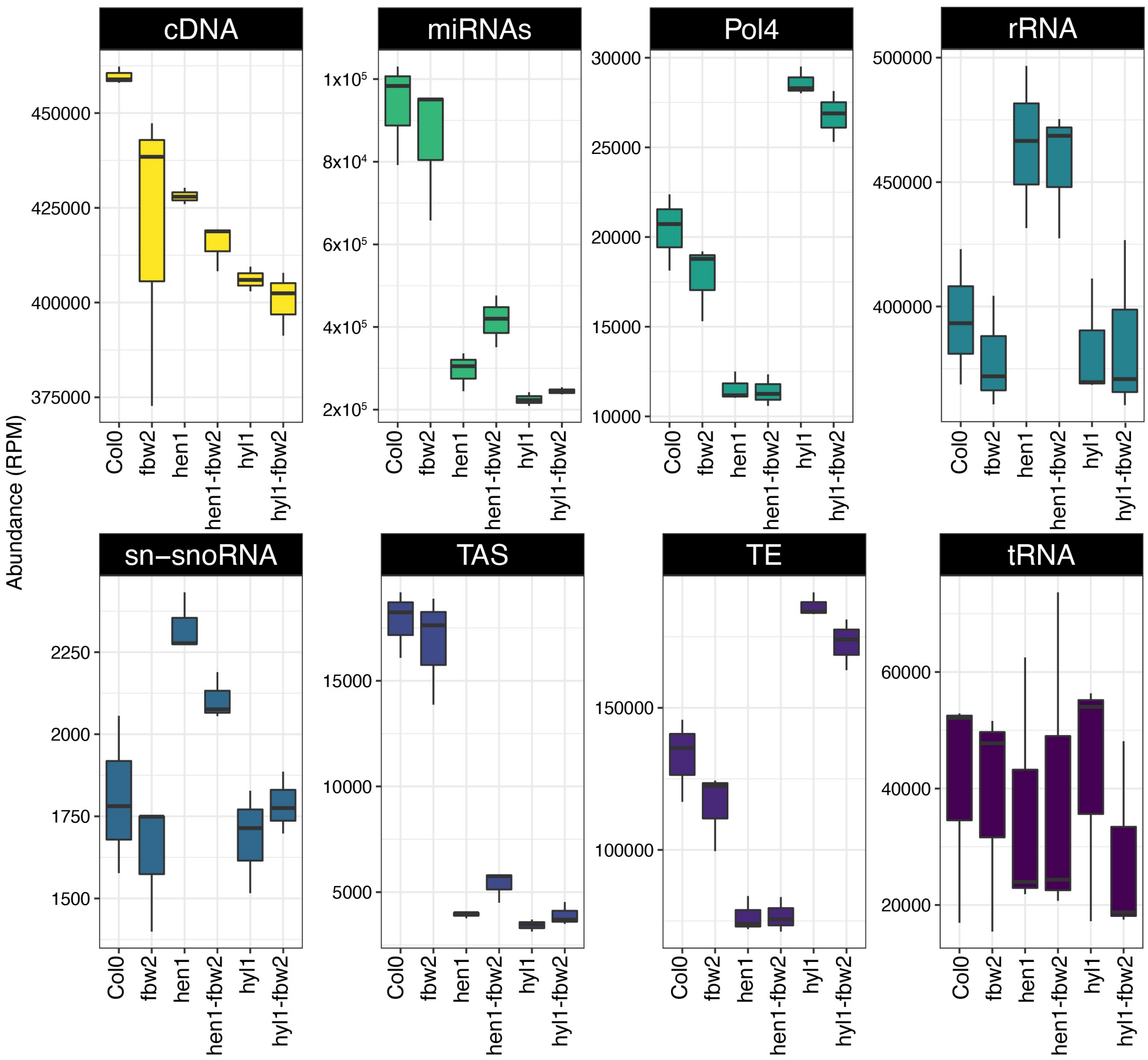


### Figure S11
